# FHF2 phosphorylation and regulation of native myocardial Na_V_1.5 channels

**DOI:** 10.1101/2023.01.31.526475

**Authors:** Adrien Lesage, Maxime Lorenzini, Sophie Burel, Marine Sarlandie, Floriane Bibault, Dan Maloney, Jonathan R. Silva, R. Reid Townsend, Jeanne M. Nerbonne, Céline Marionneau

**Affiliations:** Nantes Université, CNRS, INSERM, l’institut du thorax, F-44000 Nantes, France; Bioinformatics Solutions Inc., Waterloo, ON, Canada; Department of Biomedical Engineering, Washington University in Saint Louis, MO, USA; Departments of Cell Biology and Physiology, Washington University Medical School, Saint Louis, MO, USA; Departments of Medicine and Washington University Medical School, Saint Louis, MO, USA; Departments of Developmental Biology, Washington University Medical School, Saint Louis, MO, USA

**Keywords:** Cardiac Na_V_1.5 channels, phosphoproteomics, native FHF2 phosphorylation sites, FHF2 isoforms, neonatal and adult mouse ventricular cardiomyocytes

## Abstract

Phosphorylation of the cardiac Na_V_1.5 channel pore-forming subunit is extensive and critical in modulating channel expression and function, yet the regulation of Na_V_1.5 by phosphorylation of its accessory proteins remains elusive. Using a phosphoproteomic analysis of Na_V_ channel complexes purified from mouse left ventricles, we identified nine phosphorylation sites on Fibroblast growth factor Homologous Factor 2 (FHF2). To determine the roles of phosphosites in regulating Na_V_1.5, we developed two models from neonatal and adult mouse ventricular cardiomyocytes in which FHF2 expression is knockdown and rescued by WT, phosphosilent or phosphomimetic FHF2-VY. While the increased rates of closed-state and open-state inactivation of Na_V_ channels induced by the FHF2 knockdown are completely restored by the FHF2-VY isoform in adult cardiomyocytes, sole a partial rescue is obtained in neonatal cardiomyocytes. The FHF2 knockdown also shifts the voltage-dependence of activation towards hyperpolarized potentials in neonatal cardiomyocytes, which is not rescued by FHF2-VY. Parallel investigations showed that the FHF2-VY isoform is predominant in adult cardiomyocytes, while expression of FHF2-VY and FHF2-A is comparable in neonatal cardiomyocytes. Similar to WT FHF2-VY, however, each FHF2-VY phosphomutant restores the Na_V_ channel inactivation properties in both models, preventing identification of FHF2 phosphosite roles. FHF2 knockdown also increases the late Na^+^ current in adult cardiomyocytes, which is restored similarly by WT and phosphosilent FHF2-VY. Together, our results demonstrate that ventricular FHF2 is highly phosphorylated, implicate differential roles for FHF2 in regulating neonatal and adult mouse ventricular Na_V_1.5, and suggest that the regulation of Na_V_1.5 by FHF2 phosphorylation is highly complex.

**eTOC Summary:** Lesage *et al*. identify the phosphorylation sites of FHF2 from mouse left ventricular Na_V_1.5 channel complexes. While no roles for FHF2 phosphosites could be recognized yet, the findings demonstrate differential FHF2-dependent regulation of neonatal and adult mouse ventricular Na_V_1.5 channels.

## Introduction

Voltage-gated Na^+^ (Na_V_) channels are critical determinants of myocardial excitability, driving the fast upstroke of the action potential and conduction of electrical impulse through the myocardium (Chen-Izu et al., 2015). While most Na_V_ channels undergo rapid activation and inactivation to generate the peak Na^+^ current (I_Na_), a tiny fraction (∼0.5%) of channels remains open, leaving a small persistent Na^+^ influx, known as the late Na^+^ current (I_NaL_), which critically contributes to determining action potential duration. In ventricular cardiomyocytes, Na_V_ channels are composed primarily of the Na_V_1.5 channel pore-forming subunit, which critically functions in macromolecular protein complexes. As such, these channels are tightly embedded within local signaling domains, in which they are dynamically regulated by a rich repertoire of accessory proteins and post-translational modifications (PTMs) (Marionneau and Abriel, 2015). Defects in Na_V_1.5 channel functioning and/or regulation by these components underlie diverse forms of inherited or acquired cardiac arrhythmias. Impaired inactivation of Na_V_1.5 channels, notably, leads to alterations in channel availability and/or enhances I_NaL_, that can cause severe arrhythmias, including long QT syndrome 3, Brugada syndrome or conduction slowing. Leveraging the endogenous regulatory mechanisms of Na_V_1.5 channels is therefore essential to decipher arrhythmogenic Na_V_ current defects. Several recent studies in the laboratory demonstrated that the cardiac Na_V_1.5 protein is highly phosphorylated, and that phosphorylation-dependent regulation of Na_V_1.5 channels is critical in regulating the expression or functioning of the channel, as well as interactions with accessory proteins (Marionneau et al., 2012; Lorenzini et al., 2021). This is the case for example of serines 1933 and 1984 in the C-terminal domain of Na_V_1.5, which regulate the interaction with the Fibroblast growth factor Homologous Factor 2 (FHF2) and calmodulin, and associated channel inactivation properties (Burel et al., 2017). While numerous phosphorylation sites have been identified on the Na_V_1.5 protein, phosphorylation of the other channel complex components and the impact of these modifications on Na_V_1.5 channel expression or properties remains unappreciated.

A promising pool from which such regulation by phosphorylation may be found is the FHF accessory proteins. FHFs have emerged as pivotal players in controlling the inactivation properties of cardiac Na_V_1.5 channels, tuning both channel availability (Wang et al., 2011a; Park et al., 2016; Wang et al., 2017; Santucci et al., 2022) and I_NaL_ (Abrams et al., 2020; Gade et al., 2020; Chakouri et al., 2022). The FHF family comprises four members (FHF1=FGF12, FHF2=FGF13, FHF3=FGF11, and FHF4=FGF14), and further family diversity is achieved through the generation of several alternatively spliced isoforms highly diverging from their N-amino termini (Munoz-Sanjuan et al., 2000). FHFs are small intracellular proteins interacting directly with the membrane proximal portion of the C-terminal domain of Na_V_ channels through their common FGF homology core domain in a 1:1 stoichiometry (Goetz et al., 2009; Wang et al., 2011b). The structural basis by which FHFs regulate Na_V_ channels emerged in 2012 with the crystal structure of the ternary complex formed by a Na_V_ C-terminal domain, a FHF and Ca^2+^-free calmodulin (Wang et al., 2012), and was more recently determined by solving the cryo-electron microscopy structure of human Na_V_1.5 (Gade et al., 2020). The different FHF isoforms demonstrate species-, age-, tissue- and subcellular-specific expression patterns, and affect Na_V_ channel properties distinctively (Yang et al., 2016). In addition to their broad distribution in the nervous system, the FHF isoforms are prominently expressed in the mammalian heart. While FHF2-VY is the predominant FHF isoform in mouse ventricles, with FHF2 knockout or knockdown mice displaying severe conduction slowing (Wang et al., 2011a; Park et al., 2016; Wang et al., 2017), FHF1-B is preponderant in human hearts (Santucci et al., 2022), and has been linked to inherited arrhythmias including Brugada syndrome (Hennessey et al., 2013), long QT syndrome 3 (Liu et al., 2003), idiopathic ventricular tachycardia (Li et al., 2017), and atrial and ventricular arrhythmias with sudden cardiac death (Musa et al., 2015). In addition to its ascribed function in regulating Na_V_ channel inactivation properties, some studies have demonstrated that FHF2 also participates in regulating the surface expression of Na_V_1.5 channels in cardiomyocytes (Wang et al., 2011a; Hennessey et al., 2013; Wang et al., 2017), in a way that may be reminiscent of the well-recognized role of FHF4 in concentrating neuronal Na_V_ channels to axon initial segments and nodes of Ranvier (Goetz et al., 2009; Wang et al., 2011b).

Evidence for the role of phosphorylation in regulating FHF-dependent regulation of Na_V_ channels has mostly been provided in neuroscience studies, driven in large part by the identification of kinases that regulate the FHF-Na_V_ channel interface. Specifically, phosphorylation of neuronal Na_V_ channels by Glycogen Synthase Kinase 3 β (GSK3β) (Shavkunov et al., 2013; James et al., 2015; Hsu et al., 2017), protein kinase CK2 (Hsu et al., 2016), Ca^2+^/Calmodulin-dependent protein Kinase II (CaMKII) (Wildburger et al., 2015), as well as the tyrosine kinase Janus Kinase 2 (JAK2) (Wadsworth et al., 2020) has been demonstrated to promote binding of FHF proteins to Na_V_ channels and associated FHF-mediated channel regulation.

In this study, we investigated the pattern of phosphorylation of native mouse left ventricular FHF2 and the roles of identified FHF2 phosphorylation sites in regulating the cardiac I_Na_ current. Furthermore, by comparing the distinct consequences of FHF2 knockdown and rescue in ventricular cardiomyocytes isolated from neonatal and adult mice, as well as the FHF2 isoform expression profile, we sought to determine the differential representation and functions of FHF2 isoforms in regulating neonatal and adult mouse ventricular Na_V_ 1.5 channels.

## Results

### Identification and stoichiometry of nine native FHF2 phosphorylation sites from mouse left ventricular Na_V_ channel complexes

The identification of the FHF2 protein from mouse left ventricles was obtained from the mass spectrometric analysis of mouse left ventricular Na_V_ channel complexes purified by immunoprecipitation (IP) using an anti-Na_V_PAN mouse monoclonal (mαNa_V_PAN) antibody as previously described (Lorenzini et al., 2021). Among the 52 unique (169 total) FHF peptides detected in the mαNa_v_PAN-IPs, 38 (117 total) peptides were specific for FHF2 and conserved across the five FHF2 isoforms (**Figure 1A & Table Supplement 1**). While 8 additional unique (25 total) peptides located in the alternatively spliced N-terminus of the FHF2-VY isoform and 4 unique (24 total) peptides common to the FHF2-VY and FHF2-Y N-termini could be discriminated, no peptides specific for the three other FHF2 isoforms (FHF2-V, FHF2-A and FHF2-B) were detected. Hence, as highlighted in yellow in **Figure 1A**, 84 % of the FHF2-VY amino acid sequence was covered by mass spectrometry, representing most of the protein sequence. In addition to FHF2 peptides, sole one peptide specific for, and common to FHF1 and FHF4 sequences was identified. Altogether, these observations confirm that FHF2-VY is the predominant FHF isoform in mouse left ventricular Na_V_1.5 channel complexes, and suggest minor representations of FHF2-Y, FHF1 and/or FHF4 isoforms.

**Figure 1.**
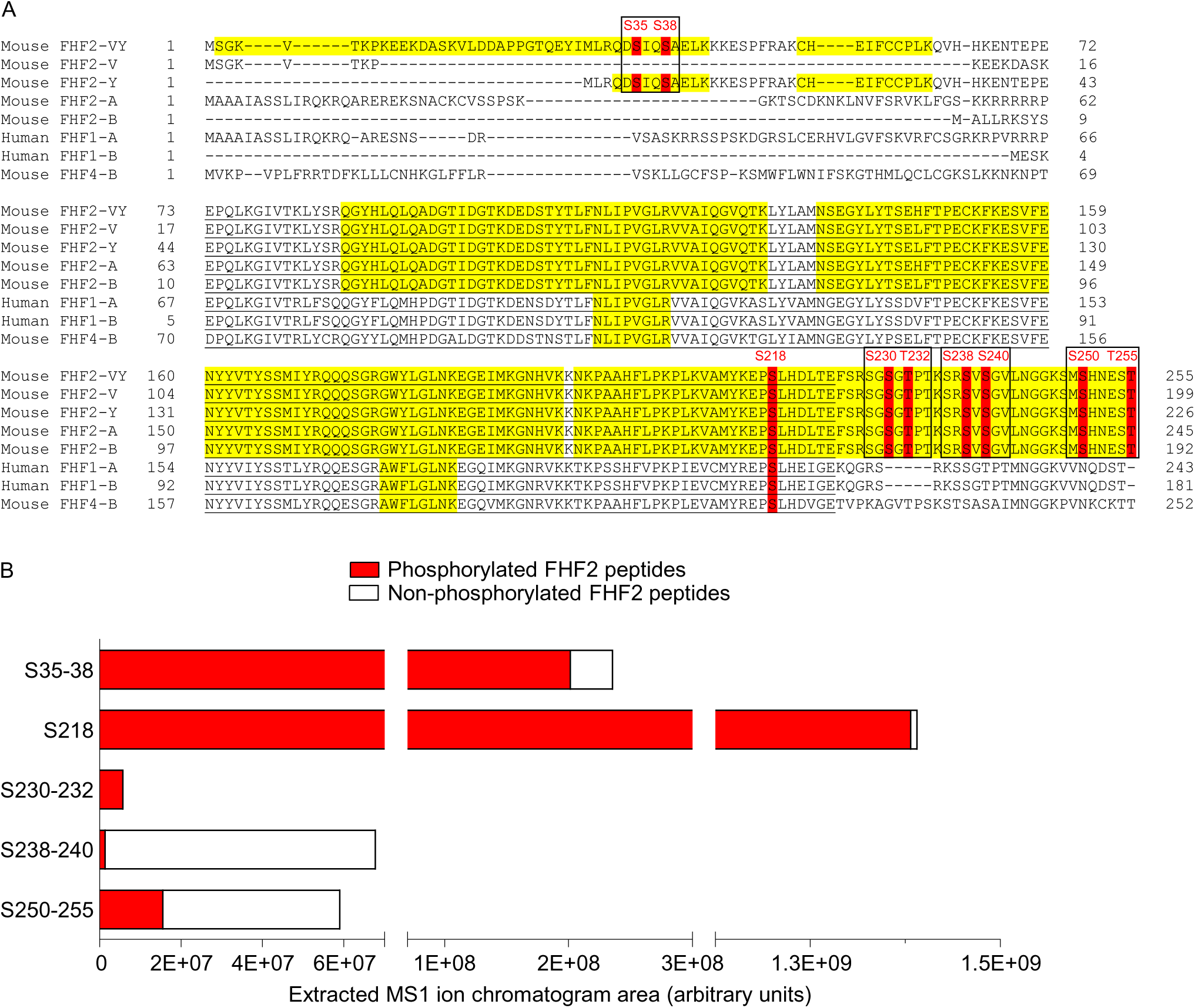
Mass spectrometric identification and stoichiometry of nine FHF2 phosphorylation sites from mouse left ventricular Na_V_ channel complexes. (**A**) The mouse FHF2-VY, FHF2-V, FHF2-Y, FHF2-A, FHF2-B and FHF4-B, and human FHF1-A and FHF1-B sequences are aligned, and the phosphorylation sites identified by MS on mouse FHF2-VY and conserved in the other FHF isoforms are highlighted in red. MS-covered sequence is highlighted in yellow; and FGF homology core domain is underlined in black. Amino acid sequences, masses and MS quality indicators of detected FHF peptides are provided in **Table 1 and Table Supplement 1**. The four phosphorylation clusters analyzed electrophysiologically are boxed in black. (**B**) The areas of extracted MS1 ion chromatograms, corresponding to MS2 spectra assigning phosphorylated (in red) and non-phosphorylated (in white) FHF2 peptides at indicated site(s), in mαNa_V_PAN-IPs from adult mouse left ventricles are indicated. Phosphosite stoichiometry is analyzed individually (S218) or by clusters of two (S35-38, S230-232, S238-240 and S250-255) as corresponding phosphosites are identified from the same phosphopeptides.

Among these 166 total FHF2 peptides, 62 peptides were phosphorylated at single or double positions, which represents more than a third of detected FHF2 peptides (**Table 1 & Table Supplement 1**). The annotation of MS/MS spectra obtained for each phosphopeptide allowed the unambiguous identification of nine phosphorylation sites on the FHF2 protein at positions S35, S38, S218, S230, T232, S238, S240, S250 and T255 (**Figure 1A**). **Table 1** lists the phosphopeptides enabling the best phosphorylation site assignment(s) for each phosphorylation site. The identification of the two N-terminal phosphorylation sites at positions S35 and S38 arises from 2 (6 total) phosphopeptides specific for FHF2-VY and 3 (16 total) phosphopeptides common to FHF2-VY and FHF2-Y, suggesting that these N-terminal phosphosites are localized on the most represented ventricular FHF2-VY isoform. Interestingly, the phosphorylation site at position S218 is conserved across all mouse FHF (1-4) isoforms, as well as the human FHF1-A and FHF1-B isoforms, while the six C-terminal FHF2 phosphorylation sites are specific for the mouse FHF2 isoforms. It is also interesting to note that, excluding S218, these phosphosites are clustered by two, which may indicate concomitant phosphorylation and involvement in a shared regulation.

**Table 1.** Phosphorylation sites, phosphopeptides and site-discriminating ions identified in FHF2 proteins from NaV channel complexes purified from adult mouse left ventricles using MS The site-discriminating ions observed in MS/MS spectra of each annotated FHF2 phosphopeptide support the assignment of the indicated phosphorylation site(s). The PEAKS Ascore is a quality indicator of site localization. The manually verified charge state of unphosphorylated and phosphorylated site-discriminating b and y ions is reported in parentheses. The (−) symbol indicates that the ion was not detected.

| Phosphorylation site(s) | Phosphopeptide sequence | m/z (charge) | AScore | b ion | Phospho b ion | y ion | Phospho y ion |
| --- | --- | --- | --- | --- | --- | --- | --- |
| S35 | 33-QD(pS)IQSAELK | 828.936 (+2) | 10 | b2 (+1) | b5 | y7 (+1) | y8 (+1) |
| S38 | 33-QDSIQ(pS)AELK | 828.933 (+2) | 38 | b5 (+1) | (-) | y4 (+1) | (-) |
| S35 + S38 | 33-QD(pS)IQ(pS)AELK | 868.917 (+2) | 1000; 1000 | b2 (+1) | b5 | y4 (+1) | y5; y8 |
| S218 | 216-EP(pS)LHDLTEFSR | 870.414 (+2) | 54 | b1 (+1) | b7 | y9 (+1) | (-) |
| S230 + T232 | 228-SG(pS)G(pT)PTKSR | 532.593 (+3) | 11; 14 | b2 | (-) | y5 (+2) | (-) |
| T232 | 228-SGSG(pT)PTKSR | 505.936 (+3) | 8 | b4 | (-) | y5 (+2) | (-) |
| S238 | 236-SR(pS)VSGVLNGGK | 566.982 (+3) | 9 | b2 | b3 (+2) | y9 | (-) |
| S240 | 238-SV(pS)GVLNGGK | 728.401 (+2) | 1000 | b2 | (-) | y7 | (-) |
| S250 | 248-SM(pS)HNEST | 609.238 (+2) | 2 | b2 | b5 (+2) | (-) | (-) |
| T255 | 248-SMSHNES(pT) | 601.240 (+2) | 12 | b7 (+2) | (-) | (-) | (-) |
| S250 + S255 | 248-SM(pS)HNES(pT) | 641.226 (+2) | 0; 0 | b2 | (-) | (-) | (-) |

Concordantly to the large number of detected phosphorylated, compared to non-phosphorylated, FHF2 peptides, further label-free quantitative analysis of the areas of extracted MS1 peptide ion chromatograms demonstrated a greater relative abundance of phosphorylated FHF2 peptides (summed area=1.6E+09 AU), compared to non-phosphorylated FHF2 peptides (summed area=1.5E+08 AU, **Figure 1B**). This analysis also revealed large differences in the relative abundance of individual FHF2 phosphopeptides. While phosphorylation at position S218 is the most abundant (area=1.4E+09 AU), followed by phosphorylation at S35-38 (area=2.0E+08 AU), phosphorylation at the six C-terminal sites at positions S250-255 (area=1.6E+07 AU), S230-232 (area=5.9E+06 AU) and S238-240 (area=1.6E+06 AU) is less represented. Noteworthy, the phosphorylated peptides assigning S35-38, S218 and S230-232 are more abundant than their non-phosphorylated counterparts, suggesting that these sites are mostly phosphorylated in mouse left ventricular Na_V_1.5 channel complexes. Taken together, these quantitative phosphoproteomic analyses identified nine phosphorylation sites on FHF2-VY from mouse left ventricular Na_V_1.5 channel complexes, among which one site at position S218 is conserved across FHF isoforms and species, and three sites at positions S35, S38 and S218 are heavily phosphorylated in mouse left ventricles.

### FHF2 knockdown and rescue in neonatal and adult mouse ventricular cardiomyocytes

In order to investigate the roles of newly-identified FHF2 phosphorylation sites in regulating the expression and/or function of the cardiac Na_V_1.5 channels, two models were developed in freshly isolated neonatal and adult mouse ventricular cardiomyocytes, and the voltage-gated Na^+^ currents were analyzed by whole-cell voltage-clamp recordings. Neonatal ventricular cardiomyocytes were isolated from wild-type (WT) mouse pups, and adult ventricular cardiomyocytes were isolated from cardiac specific FHF2-knockdown (FHF2-KD) or control FHF2-lox adult mice (Angsutararux et al., Under revision for resubmission). The knockdown of FHF2 expression in neonatal WT cardiomyocytes was obtained in culture using FHF2 shRNA-expressing adenoviruses, and was compared directly to cardiomyocytes exposed to adenoviruses expressing control shRNA. The expression of FHF2 in both neonatal and adult cardiomyocytes was then rescued using adenoviruses expressing the FHF2-VY isoform, which is the predominant FHF isoform expressed in adult mouse left ventricles (Wang et al., 2011a), in its WT, phosphosilent or phosphomimetic forms at specific site(s). Note that the human FHF2-VY cDNA sequence was used in these rescue experiments as it only differs from one amino acid (leucine 146) compared to the mouse sequence (histidine 146). With the exception of the S218 phosphosite which was mutated individually, all the other FHF2 phosphosites were mutated and analyzed by clusters of two (35­38, 230-232, 238-240 and 250-255) as indicated by the black boxes in **Figure 1A**. In the phosphosilent constructs, mutations were introduced to replace serine(s)/threonine(s) (S/T) with alanines (A), whereas in the phosphomimetic constructs, mutations were introduced to substitute glutamate(s) (E) for serine(s)/threonine(s), to mimic phosphorylation. An additional adenovirus expressing FHF2-VY phosphosilent at the nine identified sites (FHF2-VY-9A) was also generated and used as a rescue.

Quantitative RT-PCR analyses of the four FHF genes and the five FHF2 isoforms (FHF2-VY, -V, -Y, -A and -B) as well as FHF2 western blot analyses were performed to verify the specific knockdown and rescue of FHF2 in the two cardiomyocyte models. As illustrated in **Figure 2A**, the application of FHF2 shRNA-expressing adenoviruses on neonatal mouse ventricular cardiomyocytes allowed ∼90% knockdown of the transcripts encoding the FHF2-VY, FHF2-V and FHF2-A isoforms (*p*<0.0001), whereas no significant changes in the expression of FHF1, FHF2-Y and FHF4 transcripts were obtained. Consistent with previous reports (Wang et al., 2011a), the transcript expression of the FHF2-B and FHF3 isoforms were not detected in neonatal and adult mouse ventricular cardiomyocytes (data not shown). Accordingly, western blot analyses showed 99% knockdown in FHF2 protein expression (*p*<0.01) in FHF2, compared to control, shRNA-treated cardiomyocytes (**Figures 2C & 2D**). A similar knockdown in FHF2 protein expression was reached in ventricular cardiomyocytes isolated from FHF2-KD, compared to FHF2-lox, adult mice (**Figure 2F**). The expression of FHF2 was then rescued simultaneously using adenoviruses expressing the human FHF2-VY isoform in its WT, phosphosilent or phosphomimetic forms. The results from quantitative RT-PCR analyses showed that the transcript expression of the rescued human FHF2-VY constructs is in average 2- to 3-fold greater than the endogenous mouse FHF2 transcript expression (**Figure 2B**). No direct comparison of the endogenous mouse and rescued human FHF2 protein expression could be performed because the anti-FHF2 antibodies used only allowed the specific and exclusive detection of the mouse or the human FHF2 proteins (data not shown). Additionally, although the averaged rescued FHF2-VY transcript expression varied between the different adenovirus constructs, from 1.7- (for FHF2-VY-250-255E) to 5.4- (for FHF2-VY-35-38E) fold the level of endogenous FHF2-VY expression (**Figure 2B**), no significant differences in expression were observed between the WT and the different phosphosilent or phosphomimetic FHF2-VY rescued proteins (**Figure 2G**), suggesting that the observed differences in rescued transcript expression are inherent to experimental variability. Unexpectedly, however, the FHF2-VY-9A rescue demonstrated a substantial increase in transcript (8.6-fold, **Figure 2B**) and protein (3-fold, **Figure 2E**) expression, compared with the other WT or phosphomutant FHF2 adenoviral constructs, an observation that was also apparent when transfected using a plasmid vector in heterologous expression system (data not shown). Altogether, therefore, these molecular analyses validated our ability to manipulate the expression of endogenous and rescued FHF2 proteins, and thus, the possibility to examine the effects of FHF2 phosphorylation using phosphosilent or phosphomimetic FHF2-VY constructs in both neonatal and adult mouse ventricular cardiomyocytes.

**Figure 2.**
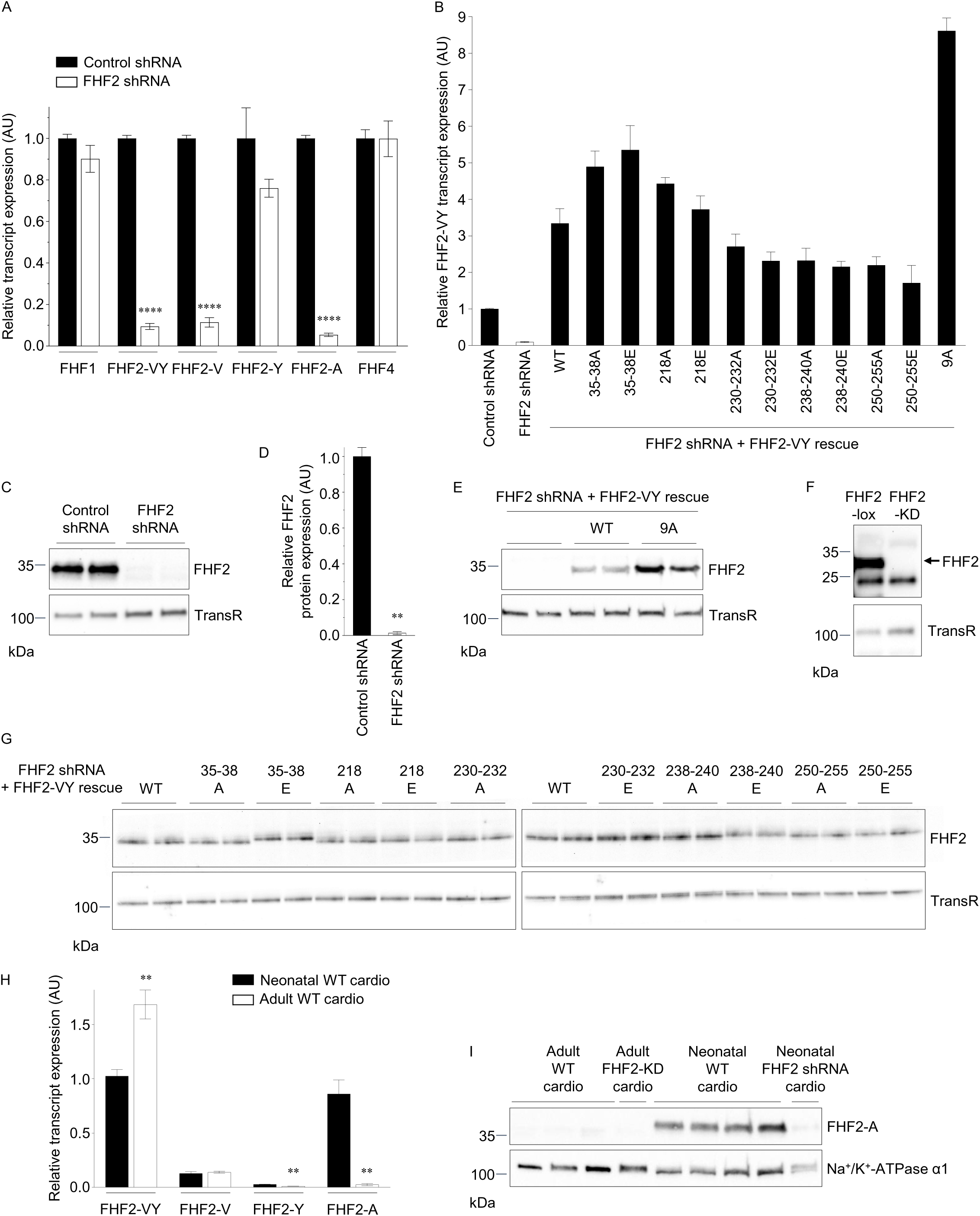
FHF2 expression in WT, knockdown and rescued neonatal and adult mouse ventricular cardiomyocytes. Neonatal ventricular cardiomyocytes were freshly isolated from WT mouse pups. Adult ventricular cardiomyocytes were freshly isolated from FHF2-lox or cardiac specific FHF2-knockdown (FHF2-KD) mice. The knockdown of FHF2 in neonatal cardiomyocytes was obtained using FHF2 shRNA-expressing adenoviruses, and the expression of FHF2 in both neonatal and adult cardiomyocytes was rescued using adenoviruses expressing WT (FHF2-VY-WT), phosphosilent (mutation to alanine) or phosphomimetic (mutation to glutamate) FHF2-VY at indicated sites. (**A**) Mean ± SEM relative transcript expression of FHF1 (n=12 in each group), FHF2-VY (n=28 in control and 24 in FHF2 shRNA samples), FHF2-V (n=12 in each group), FHF2-Y (n=4 in each group), FHF2-A (n=12 in each group) and FHF4 (n=12 in each group) isoforms in neonatal mouse ventricular cardiomyocytes infected with control or FHF2 shRNA-expressing adenoviruses. (**B**) Mean ± SEM relative transcript expression of FHF2-VY in neonatal mouse ventricular cardiomyocytes infected with adenoviruses expressing control shRNA (n=28), FHF2 shRNA alone (n=24) or with FHF2-VY-WT (n=16), FHF2-VY-35-38A (n=6), FHF2-VY-35-38E (n=4), FHF2-VY-218A (n=4), FHF2-VY-218E (n=4), FHF2-VY-230-232A (n=4), FHF2-VY-230-232E (n=4), FHF2-VY-238-240A (n=4), FHF2-VY-238-240E (n=4), FHF2-VY-250-255A (n=4), FHF2-VY-250-255E (n=4) or FHF2-VY-9A (n=2). Representative western blot (**C**) and mean ± SEM relative protein expression (**D**) of FHF2 (all isoforms) in neonatal mouse ventricular cardiomyocytes infected with adenoviruses expressing control (n=6) or FHF2 (n=6) shRNA. (**E**) Representative western blot of the rescued human FHF2-VY isoform in neonatal mouse ventricular cardiomyocytes infected with adenoviruses expressing FHF2 shRNA alone (n=4) or with FHF2-VY-WT (n=4) or FHF2-VY-9A (n=4). (**F**) Representative western blot of FHF2 (all isoforms) in ventricular cardiomyocytes isolated from FHF2-lox (n=3) and FHF2-KD (n=3) adult mice. (**G**) Representative western blots of the rescued human FHF2-VY isoform in neonatal mouse ventricular cardiomyocytes infected with adenoviruses expressing FHF2 shRNA and FHF2-VY-WT (n=14), FHF2-VY-35-38A (n=10), FHF2-VY-35-38E (n=8), FHF2-VY-218A (n=6), FHF2-VY-218E (n=4), FHF2-VY-230-232A (n=2), FHF2-VY-230-232E (n=4), FHF2-VY-238-240A (n=4), FHF2-VY-238-240E (n=4), FHF2-VY-250-255A (n=4) or FHF2-VY-250-255E (n=4). (**H**) Mean ± SEM relative transcript expression of FHF2-VY, FHF2-V, FHF2-Y and FHF2-A isoforms in neonatal and adult ventricular cardiomyocytes isolated from WT mice (n=6 in each group). (**I**) Representative western blot of FHF2-A in ventricular cardiomyocytes isolated from WT (n=6) and FHF2-KD (n=2) adult mice, and WT neonatal mouse ventricular cardiomyocytes infected (n=2) or not (n=8) with FHF2 shRNA-expressing adenoviruses. Note that the FHF2-A band is absent in neonatal cardiomyocytes knockdown for FHF2, validating the specificity of the detection. All western blots were probed in parallel with the anti-transferrin receptor (TransR) or the anti-Na^+^/K^+^-ATPase α1 antibodies to verify equal protein loading. \*\**p*<0.01, \*\*\*\**p*<0.0001 *versus* control shRNA (**A** and **D**) or neonatal WT mouse ventricular cardiomyocytes (**H**), Mann Whitney test.

### Regulation of Na_V_1.5 channels by FHF2 knockdown and rescue in neonatal mouse ventricular cardiomyocytes

The density, voltage-dependence and kinetic properties of voltage-gated Na^+^ (Na_V_) currents (I_Na_) following the knockdown and rescue of FHF2 were evaluated in neonatal mouse ventricular cardiomyocytes 48 hours after adenoviral infection using whole-cell voltage-clamp analyses. As illustrated in **Figure 3B**, and consistent with previous studies in other cardiomyocyte models (Wang et al., 2011a; Hennessey et al., 2013; Park et al., 2016; Wang et al., 2017; Santucci et al., 2022), these analyses showed that the knockdown of FHF2 significantly (*p*<0.0001) shifts the voltage-dependence of steady-state I_Na_ inactivation towards hyperpolarized potentials, compared to cardiomyocytes exposed to control shRNA-expressing adenoviruses (see distributions at -10 mV, detailed properties and statistics in **Figure 4B & Table 2**). Consistent with this effect on channel inactivation from closed state, an acceleration of the kinetics of inactivation from open state was also observed upon FHF2 knockdown (**Figures 3A & 3E**), with a significant (*p*<0.001) decrease in the time constant of fast inactivation (τ_fast_, **Figures 3F & 4D**) and an increase (*p*<0.01) in the proportion of fast to slow inactivation components (A_fast_/A_sıow_, **Figures 3H & 4F**). No significant differences in the time constant of slow inactivation (τ_slow_, **Figures 3G & 4E**), peak I_Na_ density (**Figures 3A, 3D & 4C**), time to peak I_Na_, or time for recovery from inactivation were observed upon FHF2 knockdown (**Table 2**). Interestingly, these analyses also revealed for the first time that the knockdown of FHF2 induces a significant (*p*<0.001) shift in the voltage-dependence of channel activation towards hyperpolarized potentials (**Figures 3C, 4A & Table 2**). Noteworthy, while the rescue of FHF2 expression with the WT FHF2-VY isoform partially, but significantly restored the Na_V_ channel inactivation properties from both closed (**Figures 3B, 4B & Table 2**) and open (**Figures 3A, 3E, 3F, 3H, 4D, 4F & Table 2**) states, no rescue of the activation properties was obtained (**Figures 3C, 4A & Table 2**). Together, therefore, these electrophysiological analyses suggest that an additional FHF2 isoform contributes, with FHF2-VY, to the regulation of both the activation and inactivation properties of neonatal mouse ventricular Na_V_1.5 channels.

**Table 2.**
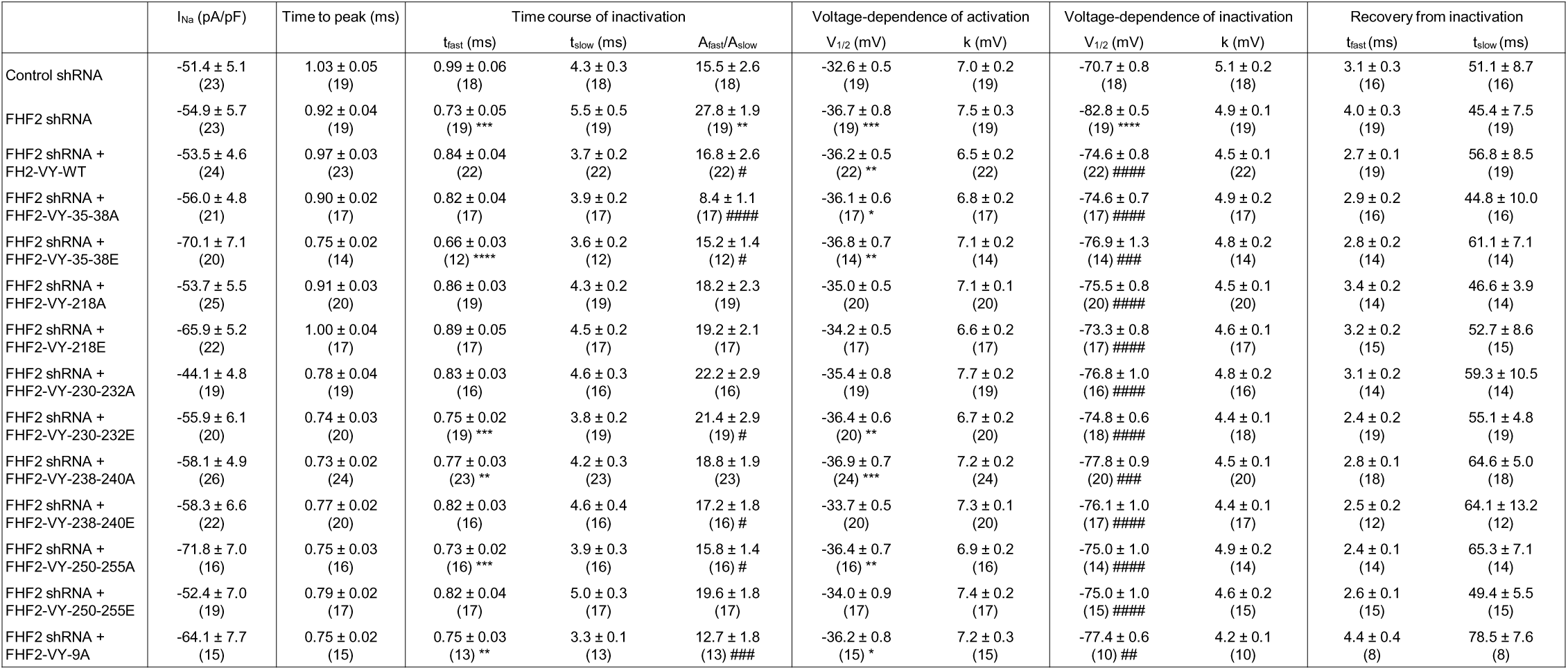
Voltage-gated Na^+^ current densities and properties in neonatal mouse ventricular cardiomyocytes infected with control shRNA, FHF2 shRNA alone or with WTor phosphomutant FHF2-VY-expressing adenoviruses Whole-cell voltage-gated Na+ currents were recorded 48 hours following infection of neonatal WT mouse ventricular cardiomyocytes with adenoviruses expressing control shRNA, FHF2 shRNA alone or with wild-type (FHF2-VY-WT), phosphosilent (mutation to alanine) or phosphomimetic (mutation to glutamate) FHF2-VY cDNA constructs using the protocols described in the materials and methods section. The peak Na+ current (I_Na_) density, time to peak I_Na_, and time course of inactivation properties presented were determined from analyses of records obtained on depolarizations to −10 mV (HP=−120 mV). All values are means ± SEM. The number of cells analyzed is provided in parentheses. \**p*<0.05, \*\**p*<0.01, \*\*\**p*<0.001, \*\*\*\**p*<0.0001 *versus* control shRNA; #*p*<0.05, ##*p*<0.01, ###*p*<0.001, ####*p*<0.0001 *versus* FHF2 shRNA; one-way ANOVA.

**Figure 3.**
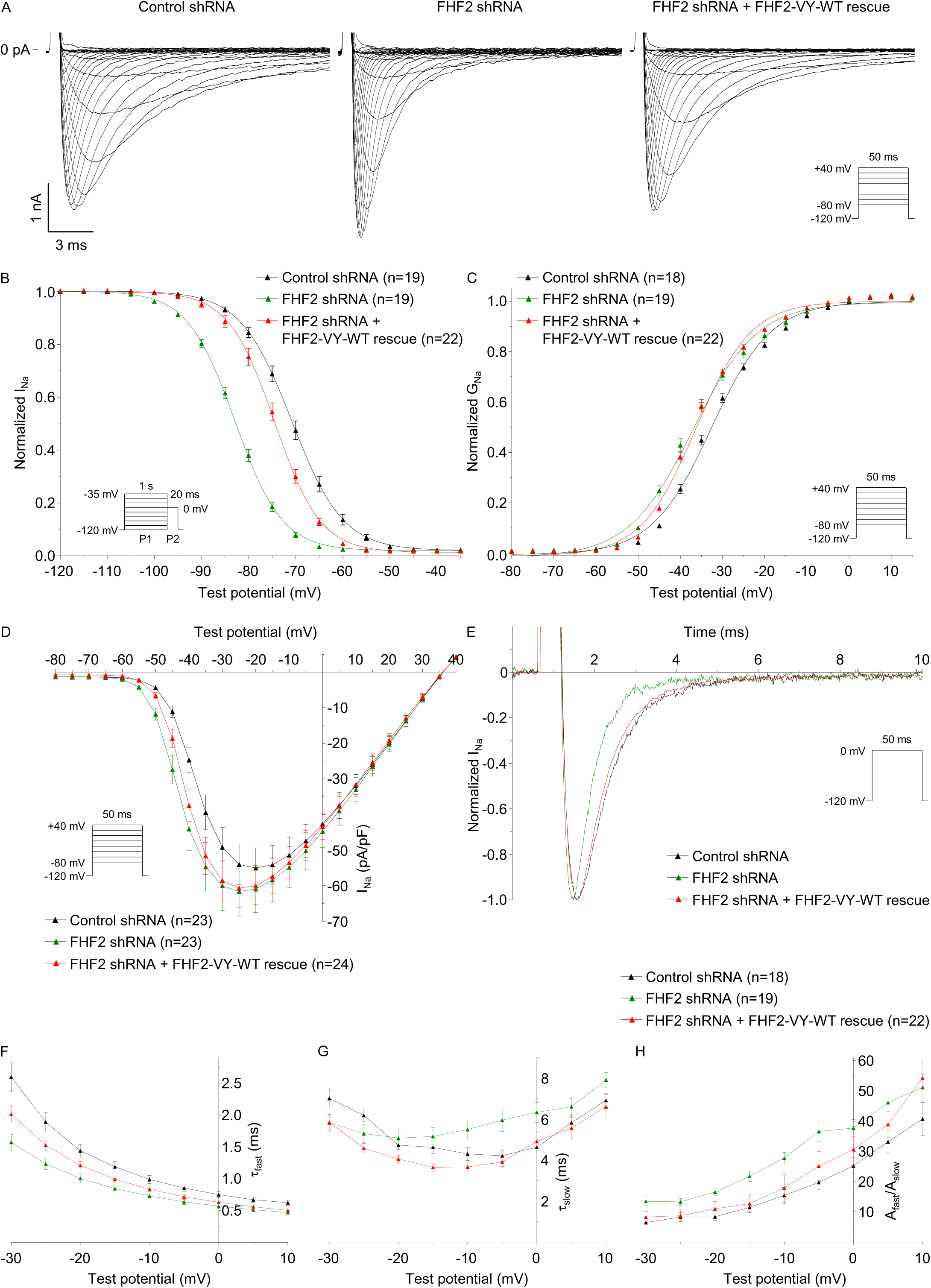
The increased closed-state and open-state inactivation rates of Na_V_ channels induced by FHF2 knockdown are partially rescued by the FHF2-VY isoform while no rescue of the shift in voltage-dependence of activation towards hyperpolarized potentials is obtained in neonatal mouse ventricular cardiomyocytes. (**A**) Representative whole-cell voltage-gated Na^+^ currents recorded 48 hours following infection of neonatal WT mouse ventricular cardiomyocytes with adenoviruses expressing control shRNA, FHF2 shRNA alone or with WT FHF2-VY (FHF2-VY-WT) using the protocols illustrated in each panel. Scale bars are 1 nA and 3 ms. Voltage-dependences of steady-state current inactivation (**B**) and activation (**C**). (**D**) Mean ± SEM peak Na^+^ current (I_Na_) densities plotted as a function of test potential. (**E**) Superimposed representative current traces recorded at 0 mV (HP=-120 mV) from cardiomyocytes infected with adenoviruses expressing control shRNA (black), FHF2 shRNA alone (green) or with FHF2-VY-WT (red). Mean ± SEM time constants of fast (τ_fast_, **F**) and slow (τ_slow_, **G**) inactivation, and proportions of fast to slow inactivation components (A_fast_/A_slow_, **H**) plotted as a function of test potential. Current densities, time- and voltage-dependent properties, as well as statistical comparisons across groups are provided in Figure 4 **& Table 2**.

**Figure 4.**
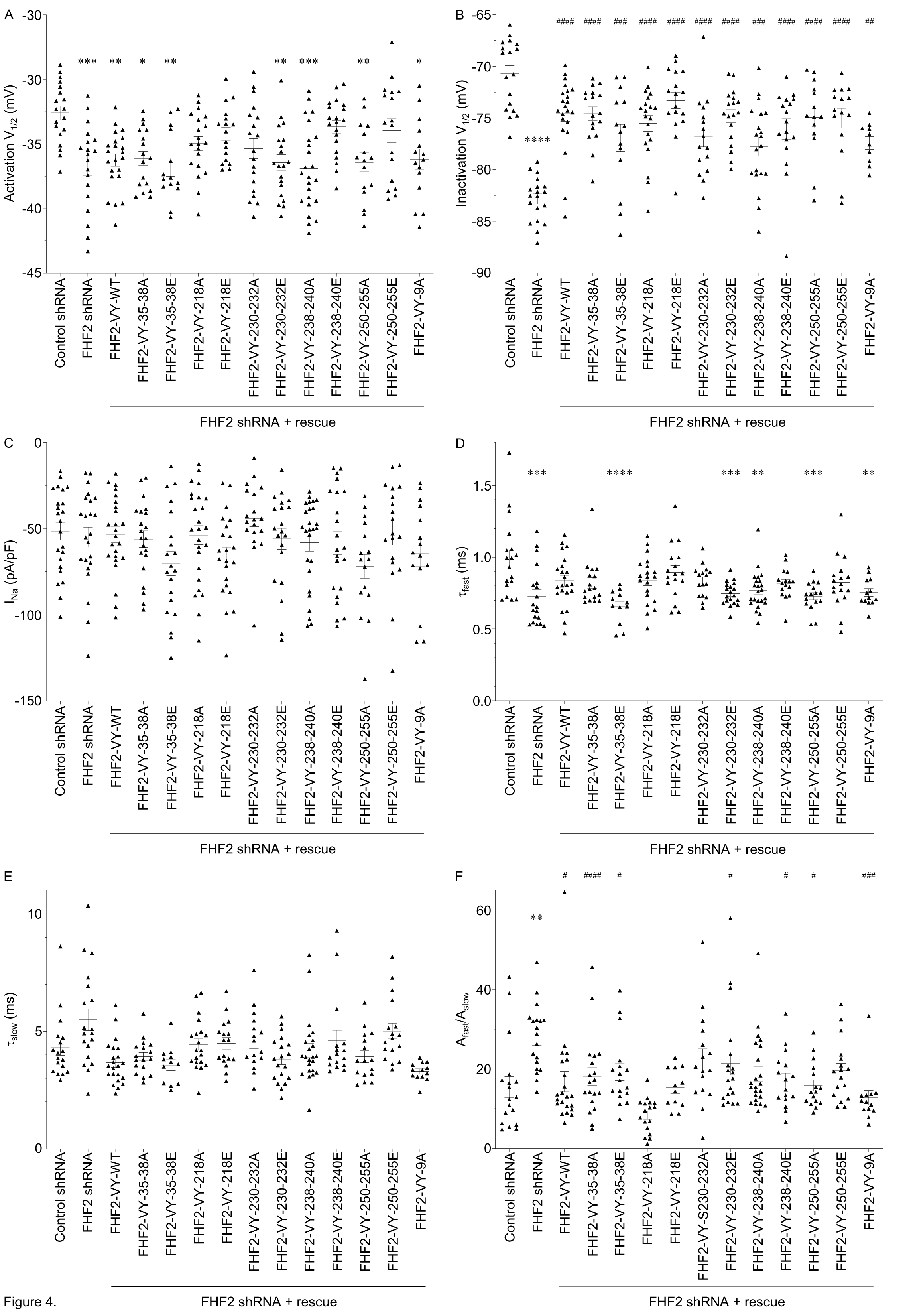
Distributions and mean ± SEM membrane potentials for half-activation (**A**) and half­inactivation (**B**), peak Na^+^ current (I_Na_) densities (**C**), time constants of fast (τ_fast_, **D**) and slow (τ_slow_, **E**) inactivation, and proportions of fast to slow inactivation components (A_fast_/A_slow_, **F**) from neonatal WT mouse ventricular cardiomyocytes infected with adenoviruses expressing control shRNA, FHF2 shRNA alone or with WT (FHF2-VY-WT), phosphosilent (mutation to alanine) or phosphomimetic (mutation to glutamate) FHF2-VY at indicated sites. Currents were recorded as described in the legend to Figure 3. The I_Na_, τ_fast_, τ_slow_ and A_fast_/A_slow_ values presented were determined from analyses of records obtained on depolarizations to -10 mV (HP=-120 mV). \**p*<0.05, \*\**p*<0.01, \*\*\**p*<0.001, \*\*\*\**p*<0.0001 *versus* control shRNA; ^#^*p*<0.05, ^##^*p*<0.01, ^###^*p*<0.001, ^####^*p*<0.0001 *versus* FHF2 shRNA; one-way ANOVA. Current densities, time- and voltage-dependent properties, as well as statistical comparisons across groups are provided in **Table 2**.

To decipher the roles of the newly-identified FHF2 phosphorylation sites in regulating the cardiac Na_V_1.5 channels, the expression of FHF2 was then rescued in neonatal mouse ventricular cardiomyocytes with the different FHF2-VY phosphomutant adenoviruses, and I_Na_ properties and densities were compared to those obtained with the WT FHF2-VY rescue. Importantly, the phosphosilent and phosphomimetic constructs for each phosphosite subgroup were compared directly to the WT FHF2-VY rescue obtained on the same days of patch-clamp analyses, and for the sake of clarity, a single representative subset of this later condition was chosen and presented in **Figure 4** and **Table 2**. To our surprise, however, no significant differences in the voltage-dependence, kinetic properties or peak I_Na_ densities were obtained for any of the ten phosphosilent or phosphomimetic FHF2-VY constructs, compared to the WT FHF2-VY rescue. A FHF2-VY construct phosphosilent at the nine identified phosphorylation sites was thus generated, but did not allow either detecting any significant changes in I_Na_ properties or density, compared to the WT FHF2-VY rescue. Together, therefore, no roles for the newly-identified FHF2 phosphorylation sites in regulating neonatal mouse ventricular Na_V_1.5 channels could be revealed, notwithstanding the multiple subgroup mutations tested. Interestingly, however, these electrophysiological analyses demonstrate for the first time that, in addition to their recognized function in regulating Na_V_1.5 channel inactivation properties, the FHF2 isoforms also regulate the voltage-dependence of Na_V_1.5 channel activation in neonatal mouse ventricular cardiomyocytes.

### Regulation of Na_V_1.5 channels by FHF2 knockdown and rescue in adult mouse ventricular cardiomyocytes

One possibility accounting for the absence of differences in the rescues of I_Na_ properties with the different phosphomutant, compared to WT, FHF2-VY constructs in neonatal mouse ventricular cardiomyocytes could arise from the fact that the FHF2 phosphorylation sites were identified from adult mouse left ventricles, which may differ from the FHF2 phosphorylation sites and/or the overall Na_V_1.5 channel complex and regulation involved in neonatal cardiomyocytes. In this respect, therefore, the same electrophysiological analyses were designed in ventricular cardiomyocytes isolated from FHF2-KD (and control FHF2-lox) adult mice (Angsutararux et al., Under revision for resubmission), 48 hours following culture and infection with the different phosphomutant (or WT) FHF2-VY adenoviruses. Similar to findings obtained in neonatal cardiomyocytes, as well as in adult cardiomyocytes isolated from the same mouse lines in the Silva laboratory (Angsutararux et al., Under revision for resubmission), the knockdown of FHF2 in adult mouse ventricular cardiomyocytes significantly increases the rate of Na_V_ channel inactivation from both closed (**Figure 5B**) and open (**Figures 5A & 5E**) states. Hence, the membrane potential of half-inactivation (V_1/2_, **Figures 5B & 6B**), the time constant of fast inactivation ^_fast_, **Figures 5F & 6D**), and the proportion of fast to slow inactivation components (A_fast_/A_slow_, **Figures 5H & 6F**) were significantly (*p*<0.0001) changed between FHF2-KD and FHF2-lox cardiomyocytes (see distributions at - 20 mV, detailed properties and statistics in **Figure 6 & Table 3**). However, contrary to findings obtained in neonatal cardiomyocytes, these analyses demonstrated the exclusive role of the FHF2-VY isoform in these effects in adult cardiomyocytes as all the Na_V_ channel inactivation properties changed with the FHF2 knockdown were completely restored by the WT FHF2-VY isoform. Additionally, the voltage-dependence of Na_V_ channel activation was not regulated by the knockdown or rescue of FHF2 expression (**Figures 5C & 6A**), which is consistent with the sole involvement of the FHF2-VY isoform in regulating Na_V_1.5 channels in these cells. Interestingly, however, the rescue of FHF2 expression with the WT FHF2-VY isoform significantly (*/*X0.0001) increases the density of the peak Na^+^ current, while no changes were observed in FHF2-KD, compared to FHF2-lox, cardiomyocytes (**Figures 5A, 5D, 6C & Table 3**).

**Table 3.**
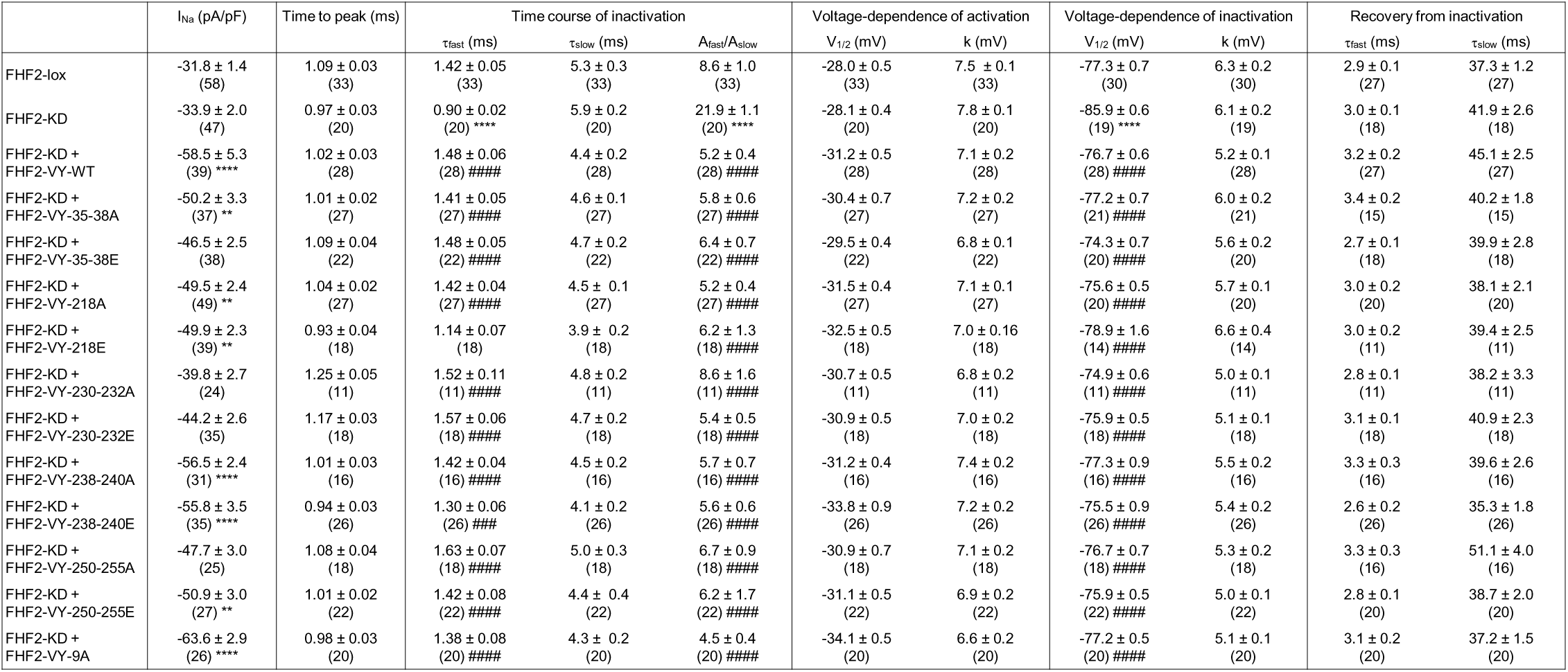
Voltage-gated Na^+^ current densities and properties in FHF2-lox and FHF2-KD adult mouse ventricular cardiomyocytes infected or not with WT or phosphomutant FHF2-VY-expressing adenoviruses Whole-cell voltage-gated Na^+^ currents were recorded 48 hours following isolation of FHF2-lox or FHF2-knockdown (FHF2-KD) adult mouse ventricular cardiomyocytes infected or not with wild-type (WT), phosphosilent (mutation to alanine) or phosphomimetic (mutation to glutamate) FHF2-VY-expressing adenoviruses using the protocols described in the materials and methods section. The peak Na^+^ current (I_Na_) density, time to peak I_Na_, and time course of I_Na_ctivation properties presented were determined from analyses of records obtained on depolarizations to −20 mV (HP=−120 mV). All values are means ± SEM. The number of cells analyzed is provided in parentheses. \**p*<0.05, \*\**p*<0.01, \*\*\**p*<0.001, \*\*\*\**p*<0.0001 *versus* FHF2-lox; #*p*<0.05, ##*p*<0.01, ###*p*<0.001, ####*p*<0.0001 *versus* FHF2-KD; one-way ANOVA.

**Figure 5.**
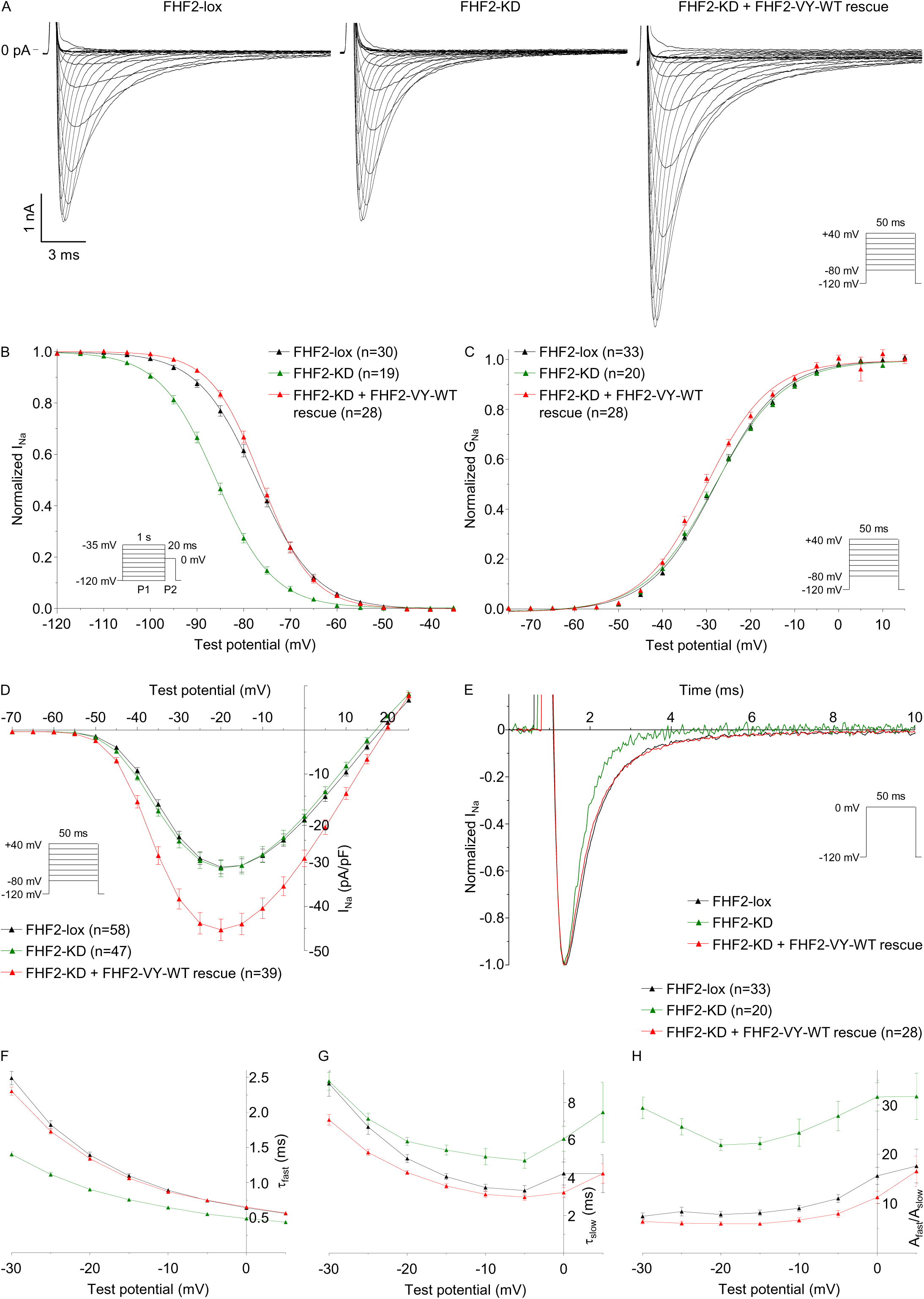
The increased closed-state and open-state inactivation rates of Na_V_ channels induced by FHF2 knockdown is completely rescued by the FHF2-VY isoform in adult mouse ventricular cardiomyocytes. (**A**) Representative whole-cell voltage-gated Na^+^ currents recorded 48 hours following isolation of FHF2-lox or cardiac specific FHF2-knockdown (FHF2-KD) adult mouse ventricular cardiomyocytes and/or infection with WT FHF2-VY (FHF2-VY-WT)-expressing adenoviruses using the protocols illustrated in each panel. Scale bars are 1 nA and 3 ms. Voltage-dependences of steady-state current inactivation (**B**) and activation (**C**). (**D**) Mean ± SEM peak Na^+^ current (I_Na_) densities plotted as a function of test potential. (**E**) Superimposed representative current traces recorded at 0 mV (HP=-120 mV) from FHF2-lox (black) or FHF2-KD adult mouse ventricular cardiomyocytes infected (red) or not (green) with FHF2-VY-WT-expressing adenoviruses. Mean ± SEM time constants of fast (τ_fast_, **F**) and slow (τ_slow_, **G**) inactivation, and proportions of fast to slow inactivation components (A_fast_/A_slow_, **H**) plotted as a function of test potential. Current densities, time- and voltage-dependent properties, as well as statistical comparisons across groups are provided in Figure 6 **& Table 3**.

**Figure 6.**
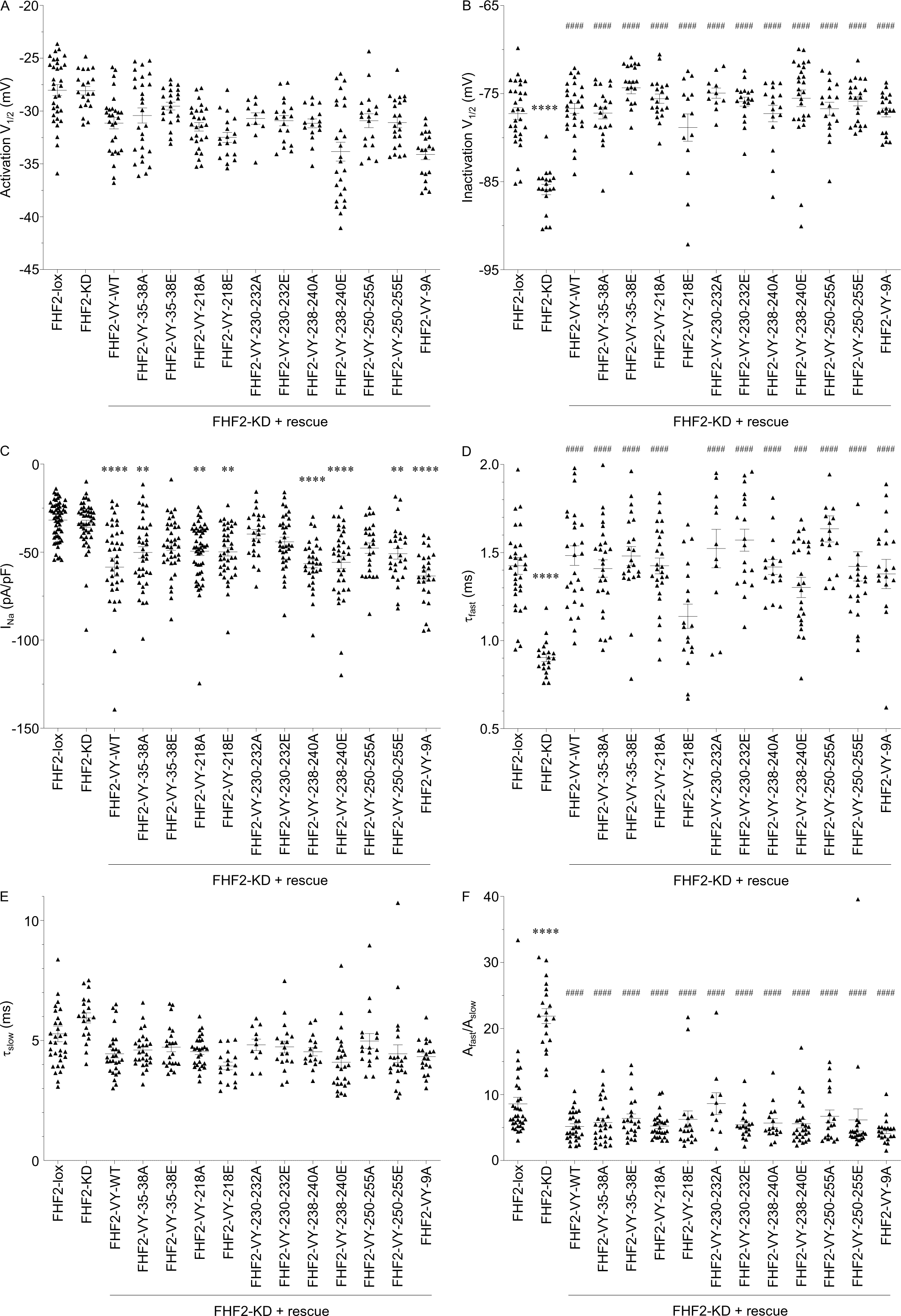
Distributions and mean ± SEM membrane potentials for half-activation (**A**) and half­inactivation (**B**), peak Na^+^ current (I_Na_) densities (**C**), time constants of fast (τ_fast_, **D**) and slow (τ_slow_, **E**) inactivation, and proportions of fast to slow inactivation components (A_fast_/A_slow_, **F**) from FHF2-lox or cardiac specific FHF2-knockdown (FHF2-KD) adult mouse ventricular cardiomyocytes infected or not with WT (FHF2-VY-WT), phosphosilent (mutation to alanine) or phosphomimetic (mutation to glutamate) FHF2-VY-expressing adenoviruses at indicated sites. Currents were recorded as described in the legend to Figure 5. The I_Na_, τ_fast_, τ_sıow_ and A_fast_/A_sıow_ values presented were determined from analyses of records obtained on depolarizations to -20 mV (HP=-120 mV). \*\**p*<0.01, \*\*\*\**p*<0.0001 *versus* FHF2-lox; ^###^*p*<0.001, ^####^*p*<0.0001 *versus* FHF2-KD; one-way ANOVA. Current densities, time- and voltage-dependent properties, as well as statistical comparisons across groups are provided in **Table 3**.

The roles of FHF2 phosphorylation sites were then explored in this adult mouse ventricular cardiomyocyte model using the different phosphosilent or phosphomimetic FHF2-VY adenoviruses. Similar to results obtained in neonatal cardiomyocytes, however, we were unable to detect any significant differences in I_Na_ densities or properties between the different phosphomutant and the WT adenoviral rescues, including with the FHF2-VY-9A rescue (**Figure 6 & Table 3**), preventing to identify any roles for FHF2 phosphorylation sites in regulating Na_V_1.5 channels.

Because it was previously shown that FHF2 also plays a crucial role in regulating the late Na^+^ current (I_NaL_) (Abrams et al., 2020; Gade et al., 2020; Chakouri et al., 2022), additional voltage-clamp experiments were designed to investigate whether the simultaneous mutation of the nine identified FHF2 phosphorylation sites to alanine, using the FHF2-VY-9A phosphomutant rescue, affects the density of TTX-sensitive I_NaL_ in adult mouse ventricular cardiomyocytes. In accordance with previous studies, these analyses demonstrated that the averaged I_NaL_ density is significantly (*/*?<0.05) increased in FHF2-KD, compared to FHF2-lox, cardiomyocytes (**Table 4**). Similar to the other Na_V_ current inactivation properties, however, comparable rescues in I_NaL_ density were obtained with the FHF2-VY-WT and FHF2-VY-9A adenoviruses.

**Table 4.**
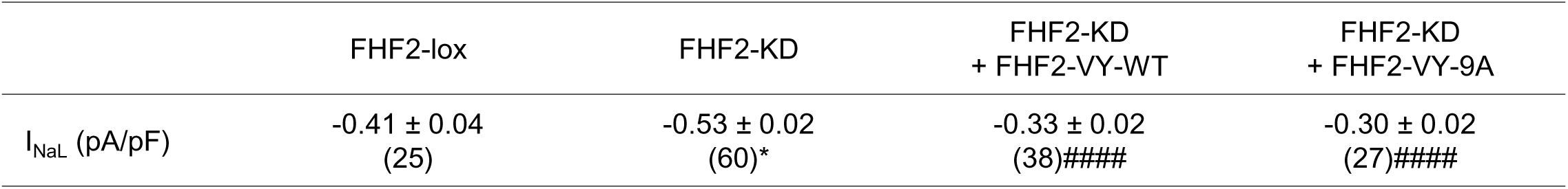
Late Na^+^ current densities in FHF2-lox and FHF2-KD adult mouse ventricular cardiomyocytes infected or not with WT or phosphosilent FHF2-VY-expressing adenoviruses The TTX-sensitive late Na^+^ current (I_NaL_) densities were measured at −20 mV (HP=−120 mV). All values are means ± SEM. The number of cells analyzed is provided in parentheses. \**p*<0.05 versus FHF2-lox; ^####^*p*<0.0001 versus FHF2-KD; one-way ANOVA.

|  | FHF2-lox | FHF2-KD | FHF2-KD<br>+ FHF2-VY-WT | FHF2-KD<br>+ FHF2-VY-9A |
| --- | --- | --- | --- | --- |
| I<sub>NaL</sub> (pA/pF) | -0.41 ± 0.04<br>(25) | -0.53 ± 0.02<br>(60)* | -0.33 ± 0.02<br>(38)#### | -0.30 ± 0.02<br>(27)#### |

Together with the electrophysiological findings in neonatal cardiomyocytes, therefore, these analyses demonstrate the exclusive role of the FHF2-VY isoform in regulating the Na_V_1.5 channel inactivation properties from closed and open states in adult mouse ventricular cardiomyocytes, and suggest the involvement of an additional FHF2 isoform in regulating both the activation and inactivation properties of Na_V_1.5 channels in neonatal mouse ventricular cardiomyocytes. Unexpectedly, however, no roles for the newly-identified FHF2 phosphorylation sites could be identified in the regulation of either neonatal or adult cardiac Na_V_1.5 channels.

### Differential transcript and protein expression of FHF2 isoforms in neonatal and adult mouse ventricular cardiomyocytes

In order to investigate the possible differences that could impart the observed distinctive effects of FHF2 knockdown and rescue on neonatal and adult ventricular Na_V_1.5 channels, further experiments were undertaken to examine the expression of the various FHF2 isoforms in freshly isolated neonatal and adult mouse ventricular cardiomyocytes. Quantitative RT-PCR analyses using isoform-specific primers demonstrated a greater (*p*<0.01) expression of the FHF2-VY isoform in adult, compared to neonatal, cardiomyocytes (**Figure 2H**). Conversely, and of potential interest in the differential regulation of neonatal Na_V_1.5 channels, a significant (*p*<0.01) higher expression of the FHF2-Y and FHF2-A isoforms was measured in neonatal, compared to adult, cardiomyocytes. Although direct comparison of relative expression of different transcripts could not reliably be achieved using the employed relative quantitative RT-PCR method, it is interesting to underscore that the FHF2-VY isoform is ∼10-fold more abundant than the three other FHF2 isoforms combined in adult cardiomyocytes, a result consistent with the exclusive mass spectrometric identification of peptides specific for the FHF2-VY (or FHF2-Y) N-terminus (**Figure 1A and Table Supplement 1**). In neonatal cardiomyocytes, however, the expression levels of the FHF2-VY and FHF2-A isoforms are similar and ∼6- to ∼30-fold greater than FHF2-V and FHF2-Y, respectively (**Figure 2H**). Remember here, nonetheless, that the FHF2-Y isoform is not of interest in the context of the mechanisms explored in the present electrophysiological analyses from neonatal cardiomyocytes as the expression of this particular isoform is not modulated by the FHF2 knockdown in these cells (**Figure 2A**). These findings thus leave the FHF2-A isoform as the sole potential FHF2 candidate responsible for the differential FHF2-dependent regulation of neonatal, compared to adult, Na_V_1.5 channels. Consistent with these findings in transcript expression, the use of an antibody specific for the FHF2-A isoform in western blot analyses demonstrated that FHF2-A is expressed in neonatal mouse ventricular cardiomyocytes, whereas no expression could be detected in adult mouse ventricular cardiomyocytes (**Figure 2I**). Together with the electrophysiological and mass spectrometric findings, therefore, these molecular analyses suggest that the FHF2-dependent regulation of Na_V_1.5 channels in adult mouse ventricular cardiomyocytes is exclusively mediated by the FHF2-VY isoform, while the FHF2-VY and FHF2-A isoforms share the regulation of neonatal Na_V_1.5 channels.

## Discussion

The results presented here provide the first phosphorylation map of the FHF2 protein isolated from native mouse left ventricular Na_V_1.5 channel complexes. Although no functional roles for the nine newly-identified FHF2 phosphorylation sites in the regulation of the expression or gating properties of Na_V_1.5-encoded channels could be recognized in the present study, the use of two distinct ventricular cardiomyocyte models from neonatal and adult mice revealed a differential FHF2-dependent regulation of Na_V_1.5 channels through the developing and adult mouse hearts. While the FHF2-VY isoform appears to be the sole FHF2 isoform involved in regulating the inactivation properties of Na_V_1.5 channels in adult mouse ventricular cardiomyocytes, our findings concur to the suggestion that both the FHF2-VY and FHF2-A isoforms share the regulation of Na_V_1.5 channel inactivation and activation properties in neonatal mouse ventricular cardiomyocytes.

### Expression and representation of FHF2 isoforms in Na_V_1.5 channel complexes in neonatal and adult mouse ventricular cardiomyocytes

Consistent with previous studies (Wang et al., 2011a), our mass spectrometric and transcript expression analyses confirmed that, out of the 4 FHF genes and the five distinct FHF2 isoforms generated by N-amino terminus alternative splicing (FHF2-VY, -V, -Y, -A and -B) reported before (Munoz-Sanjuan et al., 2000), the expression of the FHF2-VY isoform is preponderant in adult mouse ventricular cardiomyocytes, while the FHF2-B and FHF3 isoforms are not detected in either neonatal or adult mouse ventricular cardiomyocytes. Additionally, the present transcript expression findings are consistent with a differential expression of the FHF2 isoforms through the developing and adult mouse hearts, with adult mouse ventricular cardiomyocytes expressing mainly the FHF2-VY isoform, and neonatal cardiomyocytes bearing both FHF2-VY and FHF2-A. Importantly, the differential transcript expression profile of the FHF2-A isoform observed in neonatal and adult mouse ventricular cardiomyocytes was confirmed at the protein level by western blot showing specific or no FHF2-A bands in neonatal and adult samples, respectively. Altogether, therefore, these expression analyses using different technical approaches suggest that FHF2-VY is the unique FHF isoform represented in adult mouse ventricular Na_V_1.5 channel complexes, while the neonatal Na_V_1.5 channel complexes comprise both the FHF2-VY and FHF2-A isoforms.

### Proteomic and functional mapping of mouse left ventricular FHF2-VY phosphorylation sites

The present phosphoproteomic analysis confidently identified a total of nine novel native phosphorylation sites in the FHF2 protein purified from adult mouse left ventricular Na_V_1.5 channel complexes. Three of these sites, at positions S35, S38 and S218 are heavily phosphorylated in mouse left ventricles. Interestingly, the S218 phosphosite is conserved across all FHF isoforms and species, while the two N-terminal phosphosites at positions S35 and S38 are specific for FHF2-VY and FHF2-Y. Having established that the transcript expression level of the FHF2-VY isoform is predominant in adult mouse ventricular cardiomyocytes, especially compared to the much lower expression of FHF2-Y, these two N-terminal phosphosites have most likely been detected from the FHF2-VY isoform. In contrast, the six other, C-terminal FHF2 sites, at positions S230, T232, S238, S240, S250 and T255 are common to the five FHF2 isoforms, are less abundantly phosphorylated, and show a much lower stoichiometry compared to the N-terminal and S218 phosphosites. The simplest interpretation of these differences in phosphosite abundance and stoichiometry is that the first group may contribute to the basal FHF2-dependent regulatory mechanisms of cardiac Na_V_1.5 channels, while the later may participate in more local or temporary roles. While the Laezza group identified three phosphoserines on FHF4, at positions S226, S228 and S230 (Hsu et al., 2016; Hsu et al., 2017), no alignment of these three phosphoserines with the newly-identified FHF2 phosphoserines here could be obtained as the surrounding amino acid sequences are not conserved (**Figure 1A**). Additionally, while phosphorylation at Y158 was previously identified in FHF4 using *in silico* and *in vitro* analyses (Wadsworth et al., 2020), no phosphorylation was detected at the corresponding conserved FHF2 tyrosine in the present mass spectrometric analysis, likely reflecting the distinctly low level of tyrosine phosphorylation compared with phosphoserines and phosphothreonines, tissue specificity and/or differences between *in situ* and *in silico/in vitro* approaches. With the exception of the S218 phosphosite which is more isolated in the FHF2 amino acid primary sequence, the distribution of phosphosites by clusters of two led us to investigate the functional roles of these sites by four clusters of two phosphosilent or phosphomimetic mutations (35-38, 230-232, 238-240 and 250-255). To our surprise, however, no differential effects in the regulation of Na_V_1.5 channel expression or biophysical properties could be revealed with the different phosphomutant, compared to WT, FHF2-VY rescues in both neonatal and adult mouse ventricular cardiomyocyte models. A complete phosphosilent FHF2-VY construct in which the nine phosphosites were mutated to alanine (FHF2-VY-9A) was therefore generated, but did not allow either revealing any roles for FHF2 phosphorylation in regulating cardiac Na_V_1.5 channels. Although the reasons for this absence of positive findings are uncertain, especially for the FHF2-VY-9A phosphosilent mutant, three possible and non­exclusive scenarios could be incriminated. The regulation of Na_V_1.5 by phosphorylation of FHF2 could depend, as for the regulation of many protein interactions, on two distinct, direct or allosteric mechanisms. The most plausible reason, involving the more direct mechanism, may be linked to the supra-physiological levels of rescued FHF2 proteins (in average 2- to 3-fold, and 9-fold greater for FHF2-VY-9A, compared to endogenous FHF2), which may compensate for a potential loss in FHF2 interaction with the channel. Another, likely reason may be associated with the combinations of phosphosite mutations tested, which may not reflect the exact combinations of sites involved in specific channel regulations. Last, but not least, these two cardiomyocyte models, although native, may be missing some *sine qua none* molecular details involved in specific FHF2-dependent regulations, such as the activation of particular signaling pathways or kinases, therefore preventing engagement of analyzed phosphosites in specific regulatory mechanisms. Similarly, additional FHF2 phosphosites may have been lost during sample preparation and therefore not detected by mass spectrometry, which may have cancelled the occurrence of regulatory mechanisms. Although unfortunate, it is likely, however, based on the great number, abundance and/or stoichiometry of identified native FHF2 phosphosites, as well as on the previously recognized roles of phosphorylation in regulating FHF4-Na_V_ interaction and neuronal Na_V_ channel function (Shavkunov et al., 2013; James et al., 2015; Wildburger et al., 2015; Hsu et al., 2016; Hsu et al., 2017; Wadsworth et al., 2020), that the phosphorylation of FHF2 does play specific roles in regulating cardiac Na_V_1.5 channel physiology. Altogether, these notwithstanding results demonstrate that native mouse left ventricular FHF2-VY is highly phosphorylated at nine specific sites, and that the regulation of Na_V_1.5-encoded channels by FHF2 phosphorylation most certainly contributes to variable Na_V_1.5 expressivity in a highly complex manner. Further different approaches, therefore, taking the present study limitations into account, must warrant future investigations.

### FHF2 affects cardiac Na_V_ current properties and density in an age-specific manner

The study took advantage of the use of two distinct ventricular cardiomyocyte models, respectively from neonatal and adult mice, to explore the extent to which the diversity and differential expression of FHF2 isoforms may distinctly participate in the regulation of Na_V_1.5 channels through the developing and adult mouse hearts. In agreement with previous studies (Wang et al., 2011a; Park et al., 2016; Wang et al., 2017; Santucci et al., 2022), the effects of FHF2 knockdown and rescue on Na_V_ current properties herein observed demonstrate that the FHF2-VY isoform is preponderant, not to say the sole FHF2 isoform involved in regulating the Na_V_1.5 channel inactivation properties, facilitating inactivation from both closed and open states, in adult mouse ventricular cardiomyocytes. The present study also confirms the role of FHF2-VY in decreasing the late Na^+^ current in adult mouse ventricular cardiomyocytes. A differential regulation schema, however, implying not only the inactivation, but also the activation properties was observed in neonatal mouse ventricular cardiomyocytes. Based on differences in FHF2 isoform expression patterns in neonatal and adult mouse ventricular cardiomyocytes, it is tempting to speculate that both FHF2-VY and FHF2-A isoforms participate in the regulation of neonatal Na_V_1.5 channels. This conclusion, nonetheless, cannot be definitive as other differences may also exist and explain this differential regulation. It is possible, for example, that FHF2-VY does not exert the same effects on neonatal and adult Na_V_1.5 channel isoforms (Onkal et al., 2008). Alternatively, the FHF2 knockdown in neonatal cardiomyocytes may change some other channel regulatory components that are differential and therefore affect channel functioning differently compared to adult cells.

Whichever schema involved, these studies lead to the conclusion that the FHF2 isoforms are robust cellular factors in controlling the inactivation properties of Na_V_1.5 channels in both neonatal and adult mouse ventricular cardiomyocytes, whereas their role in regulating channel activation in neonatal cardiomyocytes, whether in a direct or indirect manner, would rather be secondary compared to more prevalent cellular factors. Although the diversity of the roles of the FHF2 isoforms, including FHF2-A, in regulating Na_V_ channels has previously been queried in several heterologous expression systems, as well as in dorsal root ganglion neurons or derived cells, no specific roles for FHF2-A have been attributed in the regulation of Na_V_ channel activation (Rush et al., 2006; Yang et al., 2016; Effraim et al., 2019). The most consistent finding from these studies demonstrated that FHF2-A hastens the rate of Na_V_ channel entry into the slow inactivation state and induces a dramatic slowing of recovery from inactivation, resulting in a large current decrease upon repetitive stimulations at both low and high frequencies. Together with these previous studies, therefore, the present findings suggest that this newly-identified function for FHF2-A in regulating the voltage-dependence of Na_V_ channel activation is most likely specific to the regulation of neonatal mouse ventricular Na_V_1.5 channels.

Another difference between neonatal and adult cardiomyocytes in the FHF2-dependent regulation of Na_V_1.5 channels concerns the increase in peak Na^+^ current density observed upon the FHF2 rescue in adult mouse ventricular cardiomyocytes. This finding is in a way surprising because no concordant decreased current was observed upon the FHF2 knockdown, in neither cardiomyocyte models, but is nonetheless consistent with previous studies reporting a role for FHF2 in regulating the cell surface expression of Na_V_1.5 channels (Wang et al., 2011a; Hennessey et al., 2013; Yang et al., 2016; Wang et al., 2017). In the context of the present interpretation, it is important to stress here that the expression of the rescued FHF2-VY isoforms is 2- to 3-fold greater, compared to endogenous FHF2 expression level, which may bring, by mass action, more channels to the cell surface, and therefore increase the peak Na^+^ current. Therefore, although not congruent in the literature, this novel demonstration does infer a role for FHF2-VY in the regulation of cardiac Na_V_1.5 channel cell surface expression.

In summary, our data demonstrate that ventricular FHF2 is highly phosphorylated at specific sites, and that the two main mouse ventricular FHF2 isoforms are key Na_V_1.5 channel regulatory proteins that influence membrane excitability through age- and cell environment-specific mechanisms. These novel demonstrations add to the overall suggestion that a complex and specific orchestration of regulation contributes to variable Na_V_1.5 expression and function in distinct channel complexes and contexts, and give rise to the unappreciated roles of post-translational modifications and isoform diversity in providing bases for physiological or pathological differences in Na^+^ current.

## Materials and methods

### Statement on the use of murine tissue

All investigations conformed to directive 2010/63/EU of the European Parliament, to the Guide for the Care and Use of Laboratory Animals published by the US National Institutes of Health (NIH Publication No. 85-23, revised 1985) and to local institutional guidelines.

### Immunoprecipitation of Na_V_ channel complexes

Immnunoprecipitation (IP) of Na_V_ channel complexes from mouse left ventricles was performed as described previously (Lorenzini et al., 2021). Briefly, flash-frozen left ventricles from 13-weeks-old male C57/BL6J wild-type (WT) mice were homogenized individually in ice-cold lysis buffer containing 20 mM HEPES (pH 7.4), 150 mM NaCl, 0.5% amidosulfobetaine, 1X complete protease inhibitor cocktail tablet, 1 mM phenylmethylsulfonyl fluoride (PMSF), 0.7 μg/ml pepstatin A (Thermo Fisher Scientific, Waltham, MA) and 1X Halt phosphatase inhibitor cocktail (Thermo Fisher Scientific). All reagents were from Sigma-Aldrich (Saint Louis, MO) unless otherwise noted. After 15-minutes rotation at 4°C, 8 mg of the soluble protein fractions were pre-cleared with 200 μL of protein G-magnetic Dynabeads (Thermo Fisher Scientific) for 1 hour, and subsequently used for IP with 48 μg of an anti-Na_V_PAN mouse monoclonal antibody (mαNa_V_PAN, Sigma-Aldrich, #S8809), raised against the SP19 epitope (Vassilev et al., 1988) located in the third intracellular linker loop and common to all Na_V_ channel pore-forming subunits. Prior to the IP, antibodies were cross-linked to 200 μl of protein G-magnetic Dynabeads using 20 mM dimethyl pimelimidate (Thermo Fisher Scientific) (Schneider et al., 1982). Protein samples and antibody-coupled beads were mixed for 2 hours at 4°C. Magnetic beads were then collected, washed rapidly four times with ice-cold lysis buffer, and isolated protein complexes were eluted from the beads in 1X SDS sample buffer (Bio-Rad Laboratories, Hercules, CA) at 60°C for 10 minutes. Ninety-nine percent of the immunoprecipitated mouse left ventricular Na_V_ channel protein complexes were analyzed by MS, and the remaining one percent was used to verify Na_V_1.5 IP yields by western blotting.

### Peptide preparation and isobaric labeling for LC-MS

The tryptic peptides from mouse left ventricular Na_V_ channel complexes were generated and labeled as described previously (Lorenzini et al., 2021). Briefly, the IP eluates were thawed on ice, reduced, and denatured by heating for 10 min at 95°C. The Cys residues were alkylated with iodoacetamide (10 mM) for 45 min at room temperature in the dark. The peptides were prepared using a modification (Erde et al., 2014) of the filter-aided sample preparation method (Wisniewski et al., 2009). After the addition of 300 µL of 100 mM Tris buffer (pH 8.5) containing 8 M urea (UT) and vortexing, the samples were transferred to YM-30 filter units (Millipore, MRCF0R030) and spun for 14 min at 10,000 rcf (Eppendorf, Model No. 5424). The filters were washed with 200 µl of UT buffer, and the spin-wash cycle was repeated twice. The samples were then exchanged into digest buffer with the addition of 200 µL of 50 mM Tris buffer, pH 8.0, followed by centrifugation (10,000 rcf for 10 min). After transferring the upper filter units to new collection tubes, 80 µL of digest buffer was added, and the samples were digested with trypsin (1 µg) for 4 h at 37°C. The digestion was continued overnight after adding another aliquot of trypsin. The filter units were then spun for 10 min (10,000 rcf) in an Eppendorf microcentrifuge. The filter was washed with 50 µL of Tris buffer (100 mM, pH 8.0), followed by centrifugation. The digests were extracted three times with 1 ml of ethyl acetate, and acidified to 1% trifluoroacetic acid (TFA) using a 50% aqueous solution. The pH was < 2.0 by checking with pH paper.The solid phase extraction of the peptides was performed using porous graphite carbon micro-tips (Chen et al., 2012). The peptides were eluted with 60% acetonitrile in 0.1% TFA, and pooled for drying in a Speed-Vac (Thermo Fisher Scientific, Model No. Savant DNA 120 concentrator) after adding TFA to 5%. The peptides were dissolved in 20 µL of 1% acetonitrile in water. An aliquot (10%) was removed for quantification using the Pierce Quantitative Fluorometric Peptide Assay kit (Thermo Fisher Scientific, Cat. No. 23290). The remainder of the peptides from each IP samples (∼0.5-3.5 µg) and 1.16 µg of reference pool peptide were transferred into a new 0.5 mL Eppendorf tube, dried in the Speed-Vac, and dissolved in 12 µL of HEPES buffer (100 mM, pH 8.0, Sigma-Aldrich, H3537).

The samples were labeled with tandem mass tag reagents (TMT11, Thermo Fisher Scientific) according to manufacturer’s protocol. The labeled samples were pooled, dried, and resuspended in 120 µL of 1% formic acid (FA). The TMT11 labeled sample was desalted as described above for the unlabeled peptides. The eluates were transferred to autosampler vials (Sun-Sri, Cat. No. 200046), dried, and stored at -80°C for capillary liquid chromatography interfaced to a mass spectrometer (nano-LC-MS).

### Nano-LC-MS

The mass spectrometric analysis of mouse left ventricular Na_V_ channel complexes was performed as described previously (Lorenzini et al., 2021). Briefly, the samples in formic acid (1%) were loaded (2.5 µL) onto a 75 µm i.d. × 50 cm Acclaim® PepMap 100 C18 RSLC column (Thermo Fisher Scientific) on an EASY *nano-LC* (Thermo Fisher Scientific). The column was equilibrated using constant pressure (700 bar) with 20 µL of solvent A (0.1% FA). The peptides were eluted using the following gradient program with a flow rate of 300 nL/min and using solvents A and B (acetonitrile with 0.1% FA): solvent A containing 5% B for 1 min, increased to 25% B over 87 min, to 35% B over 40 min, to 70% B in 6 min and constant 70% B for 6 min, to 95% B over 2 min and constant 95% B for 18 min. The data were acquired in data-dependent acquisition (DDA) mode. The MS1 scans were acquired with the Orbitrap™ mass analyzer over *m/z* = 375 to 1500 and resolution set to 70,000. Twelve data-dependent high-energy collisional dissociation spectra (MS2) were acquired from each MS1 scan with a mass resolving power set to 35,000, a range of *m/z* = 100 - 1500, an isolation width of 2 Th, and a normalized collision energy setting of 32%. The maximum injection time was 60 ms for parent-ion analysis and 120 ms for product­ion analysis. The ions that were selected for MS2 were dynamically excluded for 20 sec. The automatic gain control (AGC) was set at a target value of 3e6 ions for MS 1 scans and 1e5 ions for MS2. Peptide ions with charge states of one or ≥ 7 were excluded for higher-energy collision-induced dissociation (HCD) acquisition.

### MS data analysis

Peptide identification from raw MS data was performed using PEAKS Studio 8.5 (Bioinformatics Solutions Inc., Waterloo, Canada) (Zhang et al., 2012). The Uni-mouse-Reference-20131008 protein database was used for spectral matching. The precursor and product ion mass tolerances were set to 20 ppm and 0.05 Da, respectively, and the enzyme cleavage specificity was set to trypsin, with a maximum of three missed cleavages allowed. Carbamidomethylation (Cys) and TMT tags (Lys and/or peptide N-terminus) were treated as fixed modifications, while oxidation (Met), pyro-glutamination (Gln), deamidation (Asn and/or Gln), methylation (Lys and/or Arg), dimethylation (Lys and/or Arg), acetylation (Lys) and phosphorylation (Ser, Thr and/or Tyr) were considered variable modifications. The definitive annotation of each FHF2 phosphopeptide-spectrum match was obtained by manual verification and interpretation. The phosphorylation site assignments were based on the presence or absence of the unphosphorylated and phosphorylated b- and y-ions flanking the site(s) of phosphorylation, ions referred to as site-discriminating ions throughout this study. Peptide sequences, m/z, charge states, mass errors of parent ions (in ppm), PEAKS -10lgP and A scores, and charge state confirmations of site-discriminating b- and y-ions are presented in **Table Supplement 1 & Table 1**.

Label-free quantitative analysis of the areas of extracted MS1 chromatograms of phosphorylated and non-phosphorylated peptide ions covering the phosphorylation site(s) of interest was used to evaluate the proportion of phosphorylated to non-phosphorylated peptides at each position, as well as the relative abundance of phosphopeptides.

### Plasmids and adenoviruses

The FHF2 and control shRNA sequences were subcloned behind an U6 promoter into the pDUAL-U6 plasmid (Vector Biolabs, Malvern, PA). The sequence for FHF2 shRNA was 5’-CAGCACTTACACTCTGTTTAA-CTCGAG-TTAAACAGAGTGTAAGTGCTG-3’, which targets nucleotides corresponding to mouse FHF2-VY amino acids 106-113. The sequence for control shRNA was 5’-GCGCGATAGCGCTAATAATTT-CTCGAG-AAATTATTAGCGCTATCGCGC-3’, which does not correspond to any known sequence in the mouse genome. The FHF2-VY phosphomutant rescue constructs were generated by mutating the serine(s)/threonine(s) to alanine(s) (A) or glutamate(s) (E) by site-directed mutagenesis using the QuikChange II XL Site-Directed Mutagenesis kit (Agilent, Santa Clara, CA) of a pDUAL2-CCM(-) plasmid (Vector Biolabs, Malvern, PA) containing the CMV promoter in front of the human FHF2-VY cDNA (NCBI Reference Sequence NM_001139500, full-length cDNA clone purchased from Origene, Rockville, MD) silently mutated in the sequence targeted by the FHF2 shRNA. The mutated FHF2-VY constructs were then digested with restriction endonucleases to excise the mutated fragments, which were then subcloned into the original pDUAL2-CCM(-) plasmid. The pDUAL plasmids containing the shRNA or FHF2-VY constructs were then provided to Vector Biolabs (Malvern, PA) for the generation, purification and titration of recombinant (human type 5, dE1/E3) adenoviruses which also contain the Red (RFP) or Green (GFP) Fluorescent Proteins, respectively, as markers of infection, under the control of a CMV promoter. All plasmid and adenoviral constructs were sequenced to ensure that no unintentional mutations were introduced.

### Isolation, culture and adenoviral infection of neonatal mouse ventricular cardiomyocytes

Single cardiomyocytes were isolated from the ventricles of C57BL/6J WT mouse neonates aged from postnatal day 0 to 3 by enzymatic and mechanical dissociation in a semi-automated procedure by using the Neonatal Heart Dissociation kit and the GentleMACS™ dissociator (Miltenyi Biotec, Gaithersburg, MD). Briefly, hearts were harvested, and the ventricles were separated from the atria and digested in the GentleMACS™ dissociator. After termination of the program, the digestion was stopped by adding medium containing Dulbecco’s Modified Eagle’s Medium (DMEM) supplemented with 10% horse serum, 5% fetal bovine serum and 100 U/ml penicillin and 100 µg/ml streptomycin. The cell suspension was filtered to remove undissociated tissue fragments, and centrifugated. The cell pellet was resuspended in culture medium, and the cells were plated in 60 mm-diameter Petri dishes at 37°C for 1.5 hour. The non-plated cardiomyocytes were then resuspended, plated on laminin-coated dishes at a density of 5 000 or 500 000 cells per 35 mm-diameter plate for patch-clamp and molecular biology/biochemical analyses, respectively, and incubated in 37°C, 5% CO_2_: 95% air incubator. After 24 hours-plating, medium was replaced by DMEM supplemented with 1% fetal bovine serum and 100 U/mL penicillin and 100 µg/mL streptomycin, in the presence or absence of the shRNA- and FHF2-VY-expressing adenoviruses at a multiplicity of infection (MOI) of 50 and 1, respectively. Culture medium was then changed 24 and 48 hours after adenoviral infection with DMEM supplemented with 1% fetal bovine serum and 100 U/mL penicillin and 100 µg/mL streptomycin without adenoviruses.

### FHF2-lox, αMHC-Cre and FHF2-KD mice

Cardiac specific FHF2-knockdown (FHF2-KD) and control FHF2-lox adult (8-16-weeks-old) male C57BL/6J mice (Angsutararux et al., Under revision for resubmission) were used. The FHF2-lox C57BL/6J mouse line, in which the FHF2 locus is floxed, was obtained from Dr. Jeanne Nerbonne, and the αMHC-Cre C57BL/6J mouse line, expressing the Cre-recombinase driven by the cardiac specific alpha Myosin Heavy Chain (αMHC) promoter, was purchased from The Jackson Laboratory (Bar Harbor, ME, Tg(Myh6-cre)2182Mds/J mouse line). To obtain cardiac specific FHF2 targeted knockdown (FHF2-KD) mice, FHF2-lox female mice were crossed with αMHC-Cre male mice. The FHF2-lox male littermates were used as controls.

### Isolation, culture and adenoviral infection of adult mouse ventricular cardiomyocytes

Single cardiomyocytes were isolated from the ventricles of FHF2-KD and FHF2-lox adult (8-16 weeks-old) male C57BL/6J mice (Angsutararux et al., Under revision for resubmission) by enzymatic dissociation and mechanical dispersion according to a modified procedure of established methods. All reagents were from Sigma-Aldrich unless otherwise noted. Briefly, mice were injected with heparin (5000 units/kg body weight) 30 minutes before sacrifice by cervical dislocation. Hearts were quickly excised, and perfused retrogradely through the aorta with a solution at 37°C containing (in mM): NaCl, 113; KCl, 4.7; MgSO_4,_ 1.2; KH_2_PO_4_, 0.6; Nal l·IΌ.:, 0.6; HEPES, 10; NaHCO_s_, 1.6; taurine, 30; glucose, 20 (pH 7.4 with NaOH). Hearts were subsequently digested for 11 minutes with the same solution supplemented with 0.08 mg/mL Liberase TM Research Grade. Following digestion, the perfusion was stopped, the atria were removed, and the ventricles were dispersed by gentle trituration. The resulting cell suspension was filtered to remove large undissociated tissue fragments, and resuspended in solutions containing 10 mg/mL bovine serum albumin and Ca^2+^ concentrations successively increasing from nominally 0 to 0.2, 0.5 and 1 mM. Isolated cardiomyocytes were then resuspended in medium-199 supplemented with 5% fetal bovine serum, 10 mM 2,3-Butanedione monoxime, 100 U/ml penicillin and 100 µg/ml streptomycin, plated on laminin-coated dishes, and incubated in 37°C, 5% CO_2_: 95% air incubator. After 1-hour plating, culture medium was replaced by medium-199 supplemented with 0.1% bovine serum albumin, 10 mM 2,3-Butanedione monoxime, 1X Insulin/Transferrin/Sodium Selenite, 1X Chemically Defined Lipid Concentrate (Thermo Fisher Scientific), 0.5 µM cytochalasine D, 100 U/mL penicillin and 100 µg/mL streptomycin, in the presence or absence of the different FHF2-VY-expressing adenoviruses at a multiplicity of infection (MOI) of 1.

### RNA preparation and SYBR Green quantitative RT-PCR

Total RNA was isolated from cultured cardiomyocytes and analyzed using standard methods previously described in detail (Marionneau et al., 2008). Briefly, cells were washed twice in ice-cold PBS (pH 7.4) and lysed in buffer provided in the Nucleospin RNA kit (Machery-Nagel, Düren, Germany). Total RNA was isolated and DNase treated following the kit instructions. The quality of total RNA in each sample was examined by gel electrophoresis. Genomic DNA contamination was assessed by PCR amplification of each total RNA sample without prior cDNA synthesis; no genomic DNA was detected.

First strand cDNA was synthesized from 200 ng of total RNA from each sample using the High-Capacity cDNA Archive kit (Thermo Fisher Scientific). The relative expression levels of transcripts encoding the different FHF isoforms, including FHF1, FHF2-VY, FHF2-V, FHF2-Y, FHF2-A, FHF2-B, FHF3 and FHF4, as well as the hypoxanthine guanine phosphoribosyl transferase I (HPRT) used as an endogenous control, were determined by quantitative RT-PCR using 1X SYBR Green PCR Master Mix (Thermo Fisher Scientific). PCR reactions were performed on 10 ng of cDNA in the ABI PRISM 7900HT Sequence Detection System (Thermo Fisher Scientific) using isoform specific primer pairs giving 90­100% efficacy and a single amplicon at the appropriate melting temperature and size (**Table Supplement 2**). The cycling conditions included a hot start at 95°C for 10 min, followed by 40 cycles at 95°C for 15 s and 60°C for 1 min. Results for each sample were normalized to HPRT, and expressed according to the 2^- ΔCt^ method, as relative transcript expression compared with HPRT.

### Preparation of cardiomyocyte lysates and western blot analyses

Cultured cardiomyocytes were lysed and western blot analyses of cardiomyocyte lysates were completed as described previously (Lorenzini et al., 2021). Briefly, cells were washed twice in ice-cold PBS (pH 7.4) and lysed in ice-cold lysis buffer containing 20 mM HEPES (pH 7.4), 150 mM NaCl, 0.5% amidosulfobetaine, 1X complete protease inhibitor cocktail tablet, 1 mM phenylmethylsulfonyl fluoride (PMSF), 0.7 μg/ml pepstatin A (Thermo Fisher Scientific) and 1X Halt phosphatase inhibitor cocktail (Thermo Fisher Scientific). All reagents were from Sigma-Aldrich unless otherwise noted. After 15-minutes rotation at 4°C, protein concentrations in detergent-soluble cell lysates were determined using the Pierce BCA Protein Assay kit (Thermo Fisher Scientific), and proteins were subsequently analyzed by western blot. The mouse FHF2 isoforms (all included), the human FHF2-VY isoform and the mouse FHF2-A isoform were specifically detected using an anti-FHF2 rabbit polyclonal antibody (1:1000) given by Dr. Cecilia Lindskog Bergström (Human Protein Atlas project, Uppsala University, Sweden), the anti-FGF13 mouse monoclonal antibody (Antibodies Incorporated, Davis, CA, NeuroMab clone N91/27, 1:300), and the anti-Pan-FHF-A mouse monoclonal antibody (Antibodies Incorporated, NeuroMab clone N235/22, 1:300), respectively. The anti-transferrin receptor mouse monoclonal antibody (TransR, clone H68.4, Thermo Fisher Scientific, 1:1000) and the anti-alpha 1 Na^+^/K^+^-ATPase mouse monoclonal antibody (Na^+^/K^+^-ATPase α1, #ab7671, Abcam, Cambridge, United Kingdom, 1:1000) were used to verify equal protein loading. Bound primary antibodies were detected using horseradish peroxidase-conjugated goat anti-mouse or -rabbit secondary antibodies (Cell Signaling Technology, Inc., Danvers, MA), and protein signals were visualized using the SuperSignal West Dura Extended Duration Substrate (Thermo Fisher Scientific). Bands corresponding to FHF2 were normalized to bands corresponding to TransR from the same sample, and relative FHF2 protein expression is expressed relative to TransR protein expression.

### Electrophysiological recordings

Whole-cell Na_V_ currents were recorded at room temperature from neonatal and adult mouse ventricular cardiomyocytes using an Axopatch 200B amplifier (Axon Instruments, Molecular Devices, San Jose, CA) 48 hours following adenoviral infection. Voltage-clamp protocols were applied using the pClamp 10.4 software package (Axon Instruments) interfaced to the electrophysiological equipment using a Digidata 1440A digitizer (Axon Instruments). Current signals were filtered at 10 kHz prior to digitization at 50 kHz and storage. Patch-clamp pipettes were fabricated from borosilicate glass (OD: 1.5 mm, ID: 0.86 mm, Sutter Instrument, Novato, CA) using a P-97 micropipette puller (Sutter Instrument) to obtain a resistance between 0.8 and 1.5 Ml when filled with internal solution. For both neonatal and adult cardiomyocytes, the internal solution contained (in mM): NaCl 5, CsF 115, CsCl 20, HEPES 10, EGTA 10 (pH 7.35 with CsOH, ∼300 mosM). The external solution used to patch neonatal cardiomyocytes contained (in mM): NaCl 20, CsCl 103, TEA-Cl (tetraethylammonium chloride) 25, HEPES 10, Glucose 5, CaCl_2_ 1, MgCl_2_ 2, CoCl_2_ 2.5 (pH 7.4 with CsOH, ∼300 mosM); and the external solution used to patch adult cardiomyocytes contained (in mM): NaCl 10, CsCl 5, N-Methyl-D-Glucamine (NMDG) 104, TEA-Cl 25, HEPES 10, Glucose 5, CaCl_2_ 1, MgCh 2, CoCl_2_ 2.5 (pH 7.4 with CsOH, ∼300 mosM). All chemicals were purchased from Sigma-Aldrich. After establishing the whole­cell configuration, five minutes were allowed to ensure stabilization of voltage-dependence of activation and inactivation properties, at which time 25 ms voltage steps to ± 10 mV from a holding potential (HP) of -70 mV were applied to allow measurement of whole-cell membrane capacitances, input and series resistances. Only cells with access resistance < 7 MΩ were used, and input resistances were typically > 1 GΩ. After compensation of series resistance (80%), the membrane was held at a HP of -120 mV, and the voltage-clamp protocols were carried out as indicated below. Leak currents were always < 300 pA at HP (-120 mV), and were corrected offline. Cells exhibiting peak current amplitudes < 500 or > 5000 pA were excluded from analyses of biophysical properties because of errors associated with leak or voltage-clamp (Montnach et al., 2021), respectively, but were conserved in analyses of peak current density to avoid bias in evaluation of current densities.

Data were compiled and analyzed using ClampFit 11.2 (Axon Instruments), Microsoft Excel, and Prism (GraphPad Software, San Diego, CA). Whole-cell membrane capacitances (Cm) were determined by analyzing the decays of capacitive transients elicited by brief (25 ms) voltage steps to ± 10 mV from the HP (-70 mV). Input resistances were calculated from the steady-state currents elicited by the same ±10 mV steps (from the HP). Series resistances were calculated by dividing the decay time constants of the capacitive transients (fitted with single exponentials) by the Cm. To determine peak Na^+^ current-voltage relationships, currents were elicited by 50-ms depolarizing pulses to potentials ranging from -80 to +40 mV (presented at 5-s intervals in 5-mV increments) from a HP of -120 mV. Peak current amplitudes were defined as the maximal currents evoked at each voltage. Current amplitudes were leak-corrected, normalized to the Cm, and current densities are presented.

To analyze voltage-dependence of current activation properties, conductances (G) were calculated, and conductance-voltage relationships were fitted with the Boltzmann equation G = G_max_ / (1 + exp (- (V_m_ - V_1/2_) / k)), in which V_1/2_ is the membrane potential of half-activation and k is the slope factor. The time courses of inactivation of macroscopic currents were determined by fitting the current decay with the bi-exponential function I(*t*) = A_fast_ x exp (-*t*/τ_fast_) + A_slow_ x exp (-*t*/τ_slow_) + A_0_, in which A_fast_ and A_slow_ are the amplitudes of the fast and slow inactivating current components, respectively, and τ_fast_ and τ_slow_ are the decay time constants of A_fast_ and A_slow_, respectively. In order to visually inspect changes in current decay kinetics, overlays of I_Na_ recordings were obtained after normalization by the peak current amplitude; and representative current traces are presented. A standard two-pulse protocol was used to examine the voltage-dependences of steady-state inactivation. From a HP of -120 mV, 1-s conditioning pulses to potentials ranging from -120 to -35 mV (in 5-mV increments) were followed by 20-ms test depolarizations to -20 mV (interpulse intervals were 5-s). Current amplitudes evoked from each conditioning voltage were measured and normalized to the maximal current (I_max_) evoked from -120 mV, and normalized currents were plotted as a function of the conditioning voltage. The resulting steady-state inactivation curves were fitted with the Boltzmann equation I = I_max_ / (1 + exp ((V_m_ - V_1/2_) / k)), in which V_1/2_ is the membrane potential of half-inactivation and k is the slope factor. To examine the rates of recovery from inactivation, a three-pulse protocol was used. Cells were first depolarized to -20 mV (from a HP of -120 mV) to inactivate the channels, and subsequently repolarized to -120 mV for varying times (ranging from 1 to 200 ms), followed by test depolarizations to -20 mV to assess the extent of recovery (interpulse intervals were 5-s). The current amplitudes at -20 mV, measured following each recovery period, were normalized to the maximal current amplitude and plotted as function of the recovery time. The resulting plot was fitted with a double exponential function I(*t*) = A x (1 - exp (-*t* / τ_fast_) + 1 - exp (-*t* / τ_slow_)) + C to determine the time constants for fast (τ_fast_) and slow (τ_slow_) recovery from inactivation. For each of these biophysical properties, data from individual cells were first fitted and then averaged.

In experiments aimed at recording the tetrodotoxin (TTX)-sensitive late Na^+^ current (I_NaL_), cardiomyocytes were bathed in external solution containing (in mM): NaCl 120, TEA-Cl 25, HEPES 10, Glucose 5, CaCl_2_ 1, MgCl_2_ 2, CoCl_2_ 2.5 (pH 7.4 with CsOH, ∼300 mosM). Repetitive 350-ms test pulses to -20 mV from a HP of -120 mV (at 5-s intervals) were applied to cells to record Na^+^ currents in the absence of TTX. Cells were then superfused locally with the external solution supplemented with 60 μM TTX (Bio-Techne SAS, Rennes, France). Cells exhibiting differences in leak current amplitudes before and after TTX application > 5 pA at -20 mV (calculated from leak currents at -120 mV) were excluded from analyses. TTX-sensitive currents from individual cells were determined by offline digital subtraction of average leak-subtracted currents obtained from 5 recordings in the absence and in the presence of TTX after achieving steady state. The amplitude of TTX-sensitive I_NaL_ was defined as the mean steady-state current amplitude of macroscopic TTX-sensitive current measured from 150 to 350 ms. For each cell, the TTX-sensitive I_NaL_ amplitude was normalized to the Cm, and I_NaL_ current densities are presented.

### Statistical analyses

Results are expressed as means ± SEM. Data were first tested for normality using the D’Agostino and Pearson normality test. Depending on the results of normality tests, statistical analyses were then performed using the Mann-Whitney nonparametric test or the ordinary one-way ANOVA followed by the Tukey’s multiple comparisons post-hoc test, as indicated in Figures and Tables. All these analyses, as well as plots and graphs were performed using Prism (GraphPad Software).

## Acknowledgements

This work was supported by the *Agence Nationale de la Recherche* [ANR-15-CE14-0006-01 and ANR-16-CE92-0013-01 to CM] and the National Institutes of Health [R01-HL148803 to JRS and CM; R01-HL034161 and R01-HL142520 to JMN]. The proteomic experiments were performed at the Washington University Proteomics Shared Resource (WU-PSR), R. Reid Townsend MD. PhD., Director, Robert W. Sprung and Qiang Zhang, PhD., Co-Directors. The WU-PSR is supported in part by the WU Institute of Clinical and Translational Sciences (NCATS UL1 TR000448), the Mass Spectrometry Research Resource (NIGMS P41 GM103422) and the Siteman Comprehensive Cancer Center Support Grant (NCI P30 CA091842). ML was supported by a *Groupe de Réflexion sur la Recherche Cardiovasculaire-Société Française de Cardiologie* predoctoral fellowship [SFC/GRRC2018]. SB was supported by a Lefoulon-Delalande postdoctoral fellowship. The expert technical assistance of Agnès Tessier, Bérangère Evrard, Petra Erdmann-Gilmore, Dr. Yiling Mi, Alan Davis and Rose Connors is gratefully acknowledged. The content of the research reported is solely the responsibility of the authors and does not necessarily represent the official view of the funding agencies.

## Author Contributions

CM designed the study and wrote the paper. DM, RRT, JMN and CM designed, performed and/or analyzed the mass spectrometry experiments. AL, ML, SB, MS, FB, JRS, JMN and CM designed, performed and/or analyzed the functional analyses. JMN provided the FHF2-lox C57BL/6J mouse line. All authors reviewed the results and approved the final version of the manuscript.

## Abbreviations

A: alanine
E: glutamate
FHF2: Fibroblast growth factor Homologous Factor 2
FHF2-VY: isoform VY of Fibroblast growth factor Homologous Factor 2
I_Na_: peak Na^+^ current
I_NaL_: late Na^+^ current
IP: immunoprecipitation
mαNa_V_PAN: anti-Na_V_ channel subunit mouse monoclonal antibody
MS: Mass Spectrometry
MS1: mass spectrum of peptide precursors
MS2 or MS/MS: fragmentation mass spectrum of peptides selected in narrow mass range (2 Da) from MS1 scan
Na_V_: voltage-gated Na^+^ channel
pS: phosphoserine
pT: phosphothreonine
S: serine
T: threonine
WT: Wild-Type.

