## Supplemental Table 1 for "FHF2 phosphorylation and regulation of native myocardial Na_V_1.5 channels"

**Table Supplement 1.** FHF peptides identified in $Na_V$ channel complexes purified from adult mouse left ventricles using MS

| FHF isoform | Peptide sequence | m/z | Mass error (ppm) | -10lgP score |
|---|---|---|---|---|
| FHF2 | D(+229.16)EDSTYTLF | 660.314 | 0.2 | 37.7 |
| FHF2 | D(+229.16)EDSTYTLFNLIPVGLR | 1091.5841 | 0.7 | 68.6 |
| FHF2 | E(+229.16)GEIM(+15.99)K(+229.16)GNHVK(+229.16) | 487.0334 | -1.2 | 38.6 |
| FHF2 | E(+229.16)GEIMK(+229.16) | 582.8382 | -0.7 | 31.4 |
| FHF2 | E(+229.16)PS(+79.97)LHDLTEFSR | 870.4138 | 0 | 56.5 |
| FHF2 | E(+229.16)PSLHDLTEFSR | 553.9565 | 0.7 | 54.3 |
| FHF2 | E(+229.16)SVFENYYVTY | 821.8948 | -0.9 | 39.3 |
| FHF2 | E(+229.16)SVFENYYVTYSSM | 974.4464 | -1.5 | 36.4 |
| FHF2 | E(+229.16)SVFENYYVTYSSMIYR | 1190.5706 | -1.3 | 67.4 |
| FHF2 | F(+229.16)K(+229.16)ESVFENYYVTY | 1074.0575 | -1.1 | 48.8 |
| FHF2 | F(+229.16)K(+229.16)ESVFENYYVTYSSMIYR | 962.1516 | -8 | 66.2 |
| FHF2 | F(+229.16)LPK(+229.16)PLK(+229.16) | 510.6848 | 0.8 | 21.7 |
| FHF2 | G(+229.16)TIDGTK(+229.16)DEDSTYTLFNLIPVGLR | 1028.5565 | -3.7 | 52.7 |
| FHF2 | G(+229.16)WYLGLNK(+229.16) | 704.9221 | 1.1 | 43.4 |
| FHF2 | G(+229.16)WYLGLNK(+229.16)EGEIMK(+229.16) | 775.7797 | 0 | 53.1 |
| FHF2 | N(+229.16)K(+229.16)PAAHFLPK(+229.16)PLK(+229.16) | 476.3113 | 0.8 | 57.1 |
| FHF2 | N(+229.16)SEGYLYTSEHFTPEC(+57.02)K(+229.16) | 840.7415 | -0.6 | 59.4 |
| FHF2 | Q(+229.16)GYHLQLQADGTIDGTK(+229.16) | 768.4186 | 0.8 | 63.0 |
| FHF2 | Q(+229.16)GYHLQLQADGTIDGTK(+229.16)DEDSTYTLFNLIPVGLR | 1060.0593 | -0.6 | 79.0 |
| FHF2 | Q(+229.16)QQSGR | 466.7595 | 0.6 | 32.7 |
| FHF2 | S(+229.16)EGYLYTSEHFTPEC(+57.02)K(+229.16) | 802.7274 | -0.5 | 57.0 |
| FHF2 | S(+229.16)GS(+79.97)GT(+79.97)PTK(+229.16)SR | 532.5926 | 2.3 | 24.0 |
| FHF2 | S(+229.16)GSGT(+79.97)PTK(+229.16)SR | 505.9351 | -1.7 | 27.8 |
| FHF2 | S(+229.16)M(+15.99)S(+79.97)HNEST | 609.2379 | -1.8 | 30.4 |
| FHF2 | S(+229.16)MIYR | 449.7551 | 1.2 | 22.4 |
| FHF2 | S(+229.16)MS(+79.97)HNEST | 601.2413 | -0.3 | 34.7 |
| FHF2 | S(+229.16)MS(+79.97)HNEST(+79.97) | 641.2258 | 1.8 | 23.5 |
| FHF2 | S(+229.16)MSHNEST | 561.257 | -2.4 | 44.4 |
| FHF2 | S(+229.16)MSHNEST(+79.97) | 601.2401 | -2.4 | 38.7 |
| FHF2 | S(+229.16)RS(+79.97)VSGVLNGGK(+229.16) | 566.9828 | 2.1 | 36.9 |
| FHF2 | S(+229.16)SMIYR | 493.2705 | 0 | 31.0 |
| FHF2 | S(+229.16)VS(+79.97)GVLNGGK(+229.16) | 728.4011 | -1.6 | 47.8 |
| FHF2 | S(+229.16)VSGVLNGGK(+229.16) | 688.4185 | -0.8 | 55.9 |
| FHF2 | V(+229.16)AMYK(+229.16) | 535.3282 | 1 | 30.0 |
| FHF2 | V(+229.16)AMYK(+229.16)EPS(+79.97)LHDLTEF | 773.386 | -4.6 | 30.8 |
| FHF2 | V(+229.16)AMYK(+229.16)EPS(+79.97)LHDLTEFSR | 854.4339 | -0.2 | 61.0 |
| FHF2 | V(+229.16)VAIQGVQTK(+229.16) | 750.978 | -1.7 | 56.1 |
| FHF2 | Y(+229.16)K(+229.16)EPS(+79.97)LHDLTEFSR | 754.0508 | -1.6 | 51.8 |
| FHF2-VY | C(+57.02)(+229.16)HEIFC(+57.02)C(+57.02)PLK(+229.16) | 607.9788 | 0.1 | 52.5 |
| FHF2-VY | E(+229.16)EK(+229.16)DASK(+229.16) | 498.6313 | 1 | 27.9 |
| FHF2-VY | S(+229.16)GK(+42.01)VTK(+229.16)PK(+229.16)EEK(+229.16) | 548.0983 | 1.9 | 25.1 |
| FHF2-VY | V(+229.16)LDDAPPGTQEYIMLR | 1024.0393 | -0.8 | 66.3 |
| FHF2-VY | V(+229.16)LDDAPPGTQEYIMLRQDS(+79.97)IQS(+79.97)AELK(+229.16) | 884.685 | -0.4 | 53.3 |
| FHF2-VY | V(+229.16)LDDAPPGTQEYIMLRQDS(+79.97)IQSAELK(+229.16) | 864.6933 | -0.6 | 60.2 |
| FHF2-VY | V(+229.16)TK(+229.16)PK(+229.16)EEK(+229.16) | 469.5589 | 2.9 | 29.6 |
| FHF2-VY | V(+229.16)TK(+229.16)PK(+229.16)EEK(+229.16)DASK(+229.16) | 501.9194 | 2.1 | 40.8 |
| FHF2-VY/FHF2-Y | Q(+229.16)DS(+79.97)IQS(+79.97)AELK(+229.16) | 868.9174 | 0.1 | 43.5 |
| FHF2-VY/FHF2-Y | Q(+229.16)DS(+79.97)IQSAELK(+229.16) | 828.936 | 2.3 | 43.7 |
| FHF2-VY/FHF2-Y | Q(+229.16)DSIQS(+79.97)AELK(+229.16) | 828.9333 | -0.9 | 45.9 |
| FHF2-VY/FHF2-Y | Q(+229.16)DSIQSAELK(+229.16) | 788.951 | 0.1 | 55.4 |
| FHF1/FHF4 | A(+229.16)WFLGLNK(+229.16) | 703.9318 | 0.2 | 42.2 |
| FHF | N(+229.16)LIPVGLR | 555.8643 | 1.4 | 24.8 |

The peptide sequences, m/z, mass errors (in ppm) as well as the -10lgP PEAKS quality scores of unique peptides for each FHF isoform are presented. FHF2 peptides are conserved across the five FHF2 isoforms; FHF2-VY peptides are specific for FHF2-VY; FHF2-VY/FHF2-Y peptides are common to FHF2-VY and FHF2-Y; one FHF1/FHF4 peptide (second last lane) is specific for FHF1 and FHF4; and one FHF peptide (last lane) is conserved across the four FHF (1-4) proteins. Delta masses of amino acid modifications, including phosphorylation (+79.97 Da), are indicated in parentheses at specific sites in peptide sequences.
