## Supplemental Table 2 for "FHF2 phosphorylation and regulation of native myocardial Na_V_1.5 channels"

**Table Supplement 2.** Primer pairs used for SYBR Green quantitative RT-PCR

| FHF isoform | Nucleotide sequence |
|---|---|
| FHF1 | 5’-AGGCCGCGCATGGTT |
| | 5’-TCTGTTCCCCTTCATGATTTGA |
| FHF2-VY | 5‘-TTCTGCTGCCCGCTGAA |
| | 5’-GTATCCCTCGCTGTTCATTGC |
| FHF2-V | 5’-GCTTCTAAGGAGCCTCAGC |
| | 5’-CACCACCCGAAGACCCACAG |
| FHF2-Y | 5’-CGGCTCTGCGGTGTAACAG |
| | 5’-CCGAAAGGGAAATGTTTGAACA |
| FHF2-A | 5‘-CGAGAAATCCAATGCCTGC |
| | 5’-CACCACCCGAAGACCCACAG |
| FHF2-B | 5‘-GTTAAGGAAGTCATATTCAGAGC |
| | 5’-GTATCCCTCGCTGTTCATTGC |
| FHF3 | 5’-CCCGCGGTACCAAGTCACT |
| | 5’-CACCTTGGACAGCAGGATGA |
| FHF4 | 5’-CAGATGCACCCGGATGGA |
| | 5’-CAGAGTGGAATTGGTGCTGTCA |
| HPRT | 5’-TGAATCACGTTTGTGTCATTAGTGA |
| | 5‘-TTCAACTTGCGCTCATCTTAGG |
